# The multimodal action of G alpha q in coordinating growth and homeostasis in the *Drosophila* wing imaginal disc

**DOI:** 10.1101/2023.01.08.523049

**Authors:** Vijay Velagala, Dharsan K. Soundarrajan, Maria F. Unger, David Gazzo, Nilay Kumar, Jun Li, Jeremiah Zartman

**Affiliations:** Department of Chemical and Biomolecular Engineering, University of Notre Dame, Notre Dame, IN 46556; Department of Biological Sciences, University of Notre Dame, Notre Dame, IN 46556; Department of ACMS,171 Hurley Hall, University of Notre Dame, Notre Dame, IN 46556; Department of Applied and Computational Mathematics and Statistics

**Keywords:** Ca^2+^, Dilp-8, growth control, developmental timing, *Drosophila* wing disc

## Abstract

**Background:** G proteins mediate cell responses to various ligands and play key roles in organ development. Dysregulation of G-proteins or Ca^2+^ signaling impacts many human diseases and results in birth defects. However, the downstream effectors of specific G proteins in developmental regulatory networks are still poorly understood.

**Methods:** We employed the Gal4/UAS binary system to inhibit or overexpress *Gαq* in the wing disc, followed by phenotypic analysis. Immunohistochemistry and next-gen RNA sequencing identified the downstream effectors and the signaling cascades affected by the disruption of Gαq homeostasis.

**Results:** Here, we characterized how the G protein subunit Gαq tunes the size and shape of the wing in the larval and adult stages of development. Downregulation of *Gαq* in the wing disc reduced wing growth and delayed larval development. *Gαq* overexpression is sufficient to promote global Ca^2+^ waves in the wing disc with a concomitant reduction in the *Drosophila* final wing size and a delay in pupariation. The reduced wing size phenotype is further enhanced when downregulating downstream components of the core Ca^2+^ signaling toolkit, suggesting that downstream Ca^2+^ signaling partially ameliorates the reduction in wing size. In contrast, *Gαq*-mediated pupariation delay is rescued by inhibition of IP_3_R, a key regulator of Ca^2+^ signaling. This suggests that Gαq regulates developmental phenotypes through both Ca^2+^-dependent and Ca^2+^-independent mechanisms. RNA seq analysis shows that disruption of Gαq homeostasis affects nuclear hormone receptors, JAK/STAT pathway, and immune response genes. Notably, disruption of Gαq homeostasis increases expression levels of Dilp8, a key regulator of growth and pupariation timing.

**Conclusion:** Gαq activity contributes to cell size regulation and wing metamorphosis. Disruption to Gαq homeostasis in the peripheral wing disc organ delays larval development through ecdysone signaling inhibition. Overall, Gαq signaling mediates key modules of organ size regulation and epithelial homeostasis through the dual action of Ca^2+^-dependent and independent mechanisms.

## Background

How organs achieve their final size and shape during development is a fundamental question that has long puzzled biologists[1]–[4]. Additionally, the control of size and shape for multicellular systems, including synthetic organoids, is a key consideration for many applications in cell and tissue engineering[5]. Organ size control relies on both intrinsic intra-tissue and extrinsic hormonal growth regulators[6]. Information transfer between external cues and intrinsic signals is essential for maintaining robust physiology. This is achieved, in part, by G-protein coupled receptors (GPCRs), which constitute the largest family of receptors bound to the plasma membrane. GPCRs encode a diverse range of external signals into internal signaling dynamics, thereby modulating diverse physiological processes, including growth and proliferation[7]. Heterotrimeric G proteins form a vital component for the GPCR-mediated signaling cascade[8]–[10]. G proteins are coupled to GPCRs and are known to regulate a wide range of secondary messengers. In addition, G proteins transmit mitogenic signals from hormones and other ligands across the plasma membrane into the cell through its coupled receptors and mediate the cell’s response to these signals[9], [11]–[15].

G proteins are ubiquitous in plants and in all animals ranging from *Drosophila* to humans[16]. G-proteins are composed of α, β, and γ subunits that are associated with GPCRs as heterotrimeric complexes (**Fig. 1A**), where each subunit is classified into different families based on their structure and function[11]. For instance, the G*α* subunit is classified into four families: *Gα_i_*, *Gα_s_*, *Gα_12/13_*, and *Gαq*. Upon binding of the agonist, the receptor alters its conformation, leading to guanosine diphosphate (GDP) from Gα being exchanged with guanosine triphosphate (GTP)[17]. GTP-bound Gα dissociates from the heterotrimeric complex and activates phospholipase C β (PLC β)[18]. Subsequently, PLC β promotes the hydrolysis of phosphatidylinositol 4,5- biphosphate into second messengers diacylglycerol (DAG) and inositol trisphosphate (IP_3_)[19]. DAG further activates protein kinase C (PKC)[20], whereas IP_3_ stimulates the release of Ca^2+^ from the endoplasmic reticulum (ER) (**Fig. 1A, B**) by binding to the IP_3_ Receptor (IP_3_R)[19], [21]–[24]. Subsequently, Ca^2+^ binds to a range of proteins to generate diverse cellular responses[25]. More generally, G proteins transduce signals from extracellular cues to generate a range of second messengers such as IP_3_, DAG, and Ca^2+^, thus providing the framework to develop a broad range of cellular responses[26].

**Figure 1:**
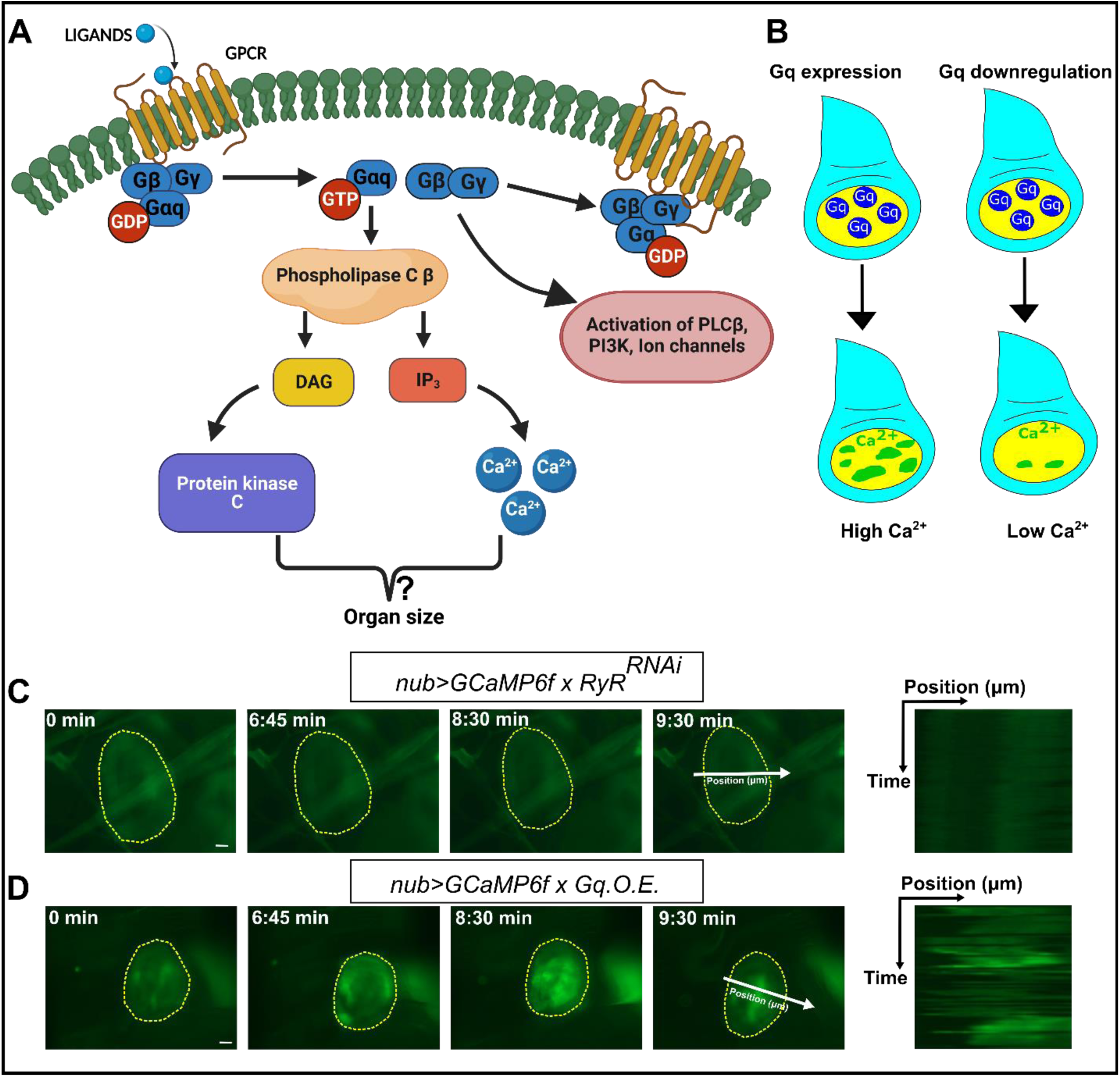
Overexpression of Gq in the wing disc pouch is sufficient to generate periodic multicellular Ca^2+^ transients and waves *in vivo*. **A)** Schematic of G Protein signaling network (partial). Hetero-trimeric proteins Gα, Gβ, and Gγ form a complex when bound to GDP. When ligands bind to G protein-coupled receptors, the Gα dissociates from the complex by exchanging GDP with GTP. GTP-bound Gαq activates the Phospholipase C β enzyme to catalyze the conversion of PIP_2_ to IP_3_ and DAG. IP_3_ binds to the IP_3_ receptor on the endoplasmic reticulum (ER), facilitating the release of stored Ca^2+^ into the cytoplasm from ER. DAG activates PKC, which further activates cAMP and other downstream signaling effectors. **B)** Schematic of a wing disc expressing wild type *Gαq* (denoted as *Gαq. O.E*) in the wing disc pouch cells using a binary expression system. High expression levels of *Gαq* result in calcium transients and waves. **C, D)**. Time-lapse images from *in vivo* imaging of wing discs expressing *RyR^RNAi^* and wildtype *Gαq* under the control of the nubbin (nub)-Gal4 driver, respectively. GCaMP6 sensor was co-expressed under the nub-Gal4 driver. Upregulation of *Gαq* in the wing disc results in the formation of periodic intercellular Ca^2+^ waves *in vivo* (D). Minimal Ca^2+^ activity, such as Ca^2+^ spikes, was observed in control discs expressing nonsense *RyR^RNAi^* (C). Yellow dotted lines indicate the wing disc pouch boundary. The UAS lines are as follows: *Gαq^WT^*, BDSC:30734, *RyR^RNAi^*, BDSC:31540. Scale bars indicate 20 µm.

Of note, Gαq serves as a key regulator of diverse biological processes, including growth, glucose transport, actin cytoskeleton regulation, cardiac physiology, and development in various model systems[13], [14], [27], [28]. For example, Gαq is essential for embryonic cardiomyocyte proliferation and cardiac development in mice[29]. Further, mice lacking both *Gαq* and *Gα11* died during embryonic development[29]. Interestingly, mice carrying a mutation for *Gαq* showed retardation in cerebellar maturation, resulting in ataxia and motor discoordination[30]. Also, the loss of *Gαq* and one *Gα11* allele in glial and precursor cells resulted in reduced somatic growth[31]. Moreover, this growth impairment is due to a decrease in the growth hormone release hormone (GHRH), thereby impairing somatotropic cell proliferation and reducing somatic growth. Surprisingly and in contrast, Parton et al. showed that a loss-of-function mouse mutant of *Gαq* in the liver and white adipose tissue results in increased body mass and hyper adiposity[27]. These genetic depletion studies highlight that the *Gαq* is essential for normal development and can act as a growth regulator, with differential phenotypes dependent on cell type. In addition to growth and development, Gαq regulates a diverse set of biological processes. For example, *Gαq* activity is required for insulin-induced glucose transport in adipocytes[13]. Further, the Adipokinetic Receptor (AkhR) functions through the Gαq/Gγ1/Plc21C signaling module to regulate body fat storage in adult *Drosophila*[28]. Mutations in the gene that encodes Gαq subunits have been shown to control skin color and hair color in mouse models[32]. Overall, these studies underscore the complexity of the diverse biological functions that Gαq performs across a broad range of biological contexts, and in many cases, the downstream pathways remain to be elucidated.

Activating *Gαq* mutations either promotes or inhibits growth based on the level of expression and cellular physiology[15]. For example, expression of the constitutive active form of *Gαq (GαqQ209L)* induced cellular transformation in NIH-3T3 cells[33], [34]. Moreover, Gαq promotes mitogenic activity through its interaction with different mitogen-activated protein kinase (MAPK) signaling pathways[35]–[39]. For instance, Garcia-Hoz et al. reported that the interaction between Gαq and PKCζ is essential for the activation of the extra-cellular-signal-regulated protein kinase (ERK) signaling cascade in cardiac myocytes and fibroblasts[40]. Additionally, activating mutations in *GNAQ,* the gene encoding the Gαq subunit in humans, contribute significantly to the progression of uveal melanoma[32], [41], [42]. However, constitutively active Gαq also can inhibit cell growth in multiple contexts[39], [43], [44]. For example, Kalinec et al. reported that the transfection of *GαqQ209L* (the mutationally active form) in NIH-3T3 cells resulted in fewer colonies and further strongly suggested that high levels of GαqQ209L are cytotoxic to cells[43]. Similar findings are reported by Qian et al., where they observed that the expression of the mutationally active form of Gαq (Gα16) partially inhibited growth in NIH-3T3 cells[45]. Likewise, the expression of an active form of Gα16 inhibits growth in small-cell lung carcinoma cells[39]. Further, this study reported an increase in c-Jun-N-terminal-Kinase (JNK) signaling cascade upon Gα16 expression, thereby suggesting a possible mechanism for growth inhibition. Based on these contrasting effects, it can be inferred that the mutationally active form of Gαq expression regulates growth in a cell-specific manner and subsequently induce different growth phenotypes. However, despite advances in characterizing the functional roles of G proteins in a range of cell types, there is a lack of a general systems-level understanding of how G protein signaling impacts organ growth.

Previously, we have shown that perturbing *Gαq* expression regulates the size of *Drosophila* wings [46]. Further, we identified that the overexpression of wildtype *Gαq* is correlated with the occurrence of intercellular Ca^2+^ waves (ICWs) in *ex vivo* wing disc cultures[46]. Further, Gαq is required for wound-induced Ca^2+^ waves in the pupal notum[47], [48]. Cumulatively, Gαq can either promote or inhibit growth depending on the cellular context and the expression levels. However, the precise molecular mechanisms through which Gαq transduces the growth regulatory signals are not known. This can be primarily attributed due to the wide range of second messengers it activates. For example, the contribution of Gαq-mediated Ca^2+^ signaling toward regulating organ growth parameters, including size and shape, is currently unknown.

To address this question in a well-defined and genetically accessible system, we used the *Drosophila* wing disc to identify the key downstream genes regulated downstream of *Gαq* loss- of-function and gain-of-function perturbations. The *Drosophila* wing imaginal disc is a key model for studying epithelial morphogenesis and signal transduction pathways, including Ca^2+^ signaling[2], [49]–[52]. With the plethora of available genetic tools to generate organ or cell-specific mutants along with the short lifecycle of the *Drosophila*, the wing disc system provides a powerful platform to decode complex signaling mechanisms[53]. The wing disc is invaluable for phenotypic screening in early preclinical studies[54]–[56]. Multiple signaling pathways, such as the Bone Morphogenetic Protein BMP (*Drosophila* Decapentaplegic/Dpp), Wnt (*Drosophila* Wg), and Hedgehog (Hh), coordinate the patterning of epithelial cells and were discovered in the wing imaginal disc[57]–[59]. In *Drosophila*, Gαq plays a key role in regulating neuronal pathfinding[60]. Additionally, it is involved in regulating gut innate immunity through the modulation of DUOX[61]. Also of note, GPCRs are known to regulate metamorphosis by transmitting ecdysone signaling and controlling hormone production through the nuclear hormone pathway[62]–[66]. Altogether, these studies have established different functional roles assumed by Gαq in *Drosophila*. However, the organ-specific role of Gαq in the growth regulation of peripheral non-neural organ development is relatively less studied.

Here, we combined genetic, phenotypic, and transcriptomic analysis to map the phenotypic consequences of perturbing Gαq homeostasis during organ development to downstream biological processes and pathways. Organ size regulation extensively relies on robust coordination between growth rate and developmental time. We show that *Gαq* perturbations affect both the wing growth and the larval-to-pupal transition time, thereby affecting the final wing size. Our RNA-seq analysis confirms that Gαq modulates key signaling events that regulate growth and developmental time. Thus, Gαq orchestrates the final wing size by interacting with key components of the signaling cascades regulating growth and developmental timing. Additionally, we investigated whether suppressing Gαq-mediated calcium signaling suppresses the reported phenotypes. We found that suppressing calcium signaling components downstream of *Gαq* overexpression suppressed the pupariation delay and further decreased growth. These results suggest that the balance of Gαq regulation of growth through Ca^2+^-dependent and Ca^2+^- independent mechanisms determine the net phenotypic consequences of perturbed Gαq activity during wing development. Overall, this study highlights the multimodal action of the Gαq in balancing growth through the dual action of its protein-protein interactions and secondary messenger regulation and provides a potential mechanism that explains the differential growth outcomes across biological contexts.

## Results

### Gαq homeostasis is necessary for optimal organ size

Gαq signaling has been demonstrated to be essential for the initiation of Ca^2+^ waves in wounded tissues[48]. In addition, our previous study showed that *Gαq* overexpression leads to the formation of intracellular calcium waves (ICWs) in the wing disc pouch cells cultured *ex vivo*[25], [46]. Here, we confirmed this result *in vivo* by imaging larva co-expressing both GCaMP6f and *Gαq* in the wing disc (**Fig. 1C, D**). Both *ex vivo* and *in vivo* imaging confirmed that the *Gαq* expression in the wing disc leads to the formation of regular and repeating ICWs.

Either overexpression or RNAi-mediated inhibition of *Gαq* in the wing disc pouch cells leads to reduced adult wing size, but it is unknown which downstream growth regulatory pathways might be impacted[46], [67]. To further characterize the roles of Gαq in regulating organ growth, we extended and confirmed our phenotypic analysis by testing the impact of *Gαq* perturbation using additional Gal4 drivers, including nubbin-Gal4, MS1096-Gal4, and C765-Gal4. The reported experiments utilize a binary expression system based on the expression of the yeast Gal4 transcription factor, which is driven by a tissue-specific promoter[53]. Consequently, analysis with multiple Gal4 drivers helps to compare the impact of a given Gal4 line. We next quantified the phenotypic consequences for both overexpression and knockdown of Gαq levels. First, we measured the average wing size for the control crosses of each Gal4 line expressing *RyR^RNAi^*, a control construct that has no known phenotype and targets a gene with no known expression in the wing disc[67] (**Fig. 2A-C**). Next, we knocked down *Gαq* with RNAi constructs using multiple Gal4 drivers (**Fig. 2D-F**). Interestingly, *Gαq* knockdown under the control of the nubbin-Gal4 driver leads to a failure in wing expansion (**Fig. 2D**). However, the other Gal4 drivers led to relatively normally patterned wings with reduced wing size (**Fig. 2E, F, J**). This phenotypic variance for different Gal4 drivers may be attributed to the variation in the spatial-temporal expression patterns of the Gal4 drivers. For instance, MS1096-Gal4 is expressed strongly in the dorsal region of the wing disc pouch during larval and prepupal stages[68], [69], whereas the C765-Gal4 driver is uniformly active in the larval wing disc from the second instar onwards[70]. The protein product of the *nubbin* gene is present from the early second instar onwards but strongly expressed in the third instar discs[71]. Next, we overexpressed *Gαq* in the wing disc using the Gal4 drivers mentioned above. Interestingly, we observed a reduction in the wing size for all the Gal4 drivers. Overexpression of *Gαq* with the MS1096-Gal4 driver showed a smaller wing size when compared with nubbin-Gal4 and C765-Gal4 drivers (**Fig. 2J**). Thus, these results indicate that Gαq homeostasis is essential for normal wing disc growth. Notably, either downregulation or upregulation reduces the final wing size. Further, the severity of the phenotype associated with *Gαq* downregulation varies depending on the Gal4 driver used for the transgenic expression of Gαq*_RNAi_*.

**Figure 2:**
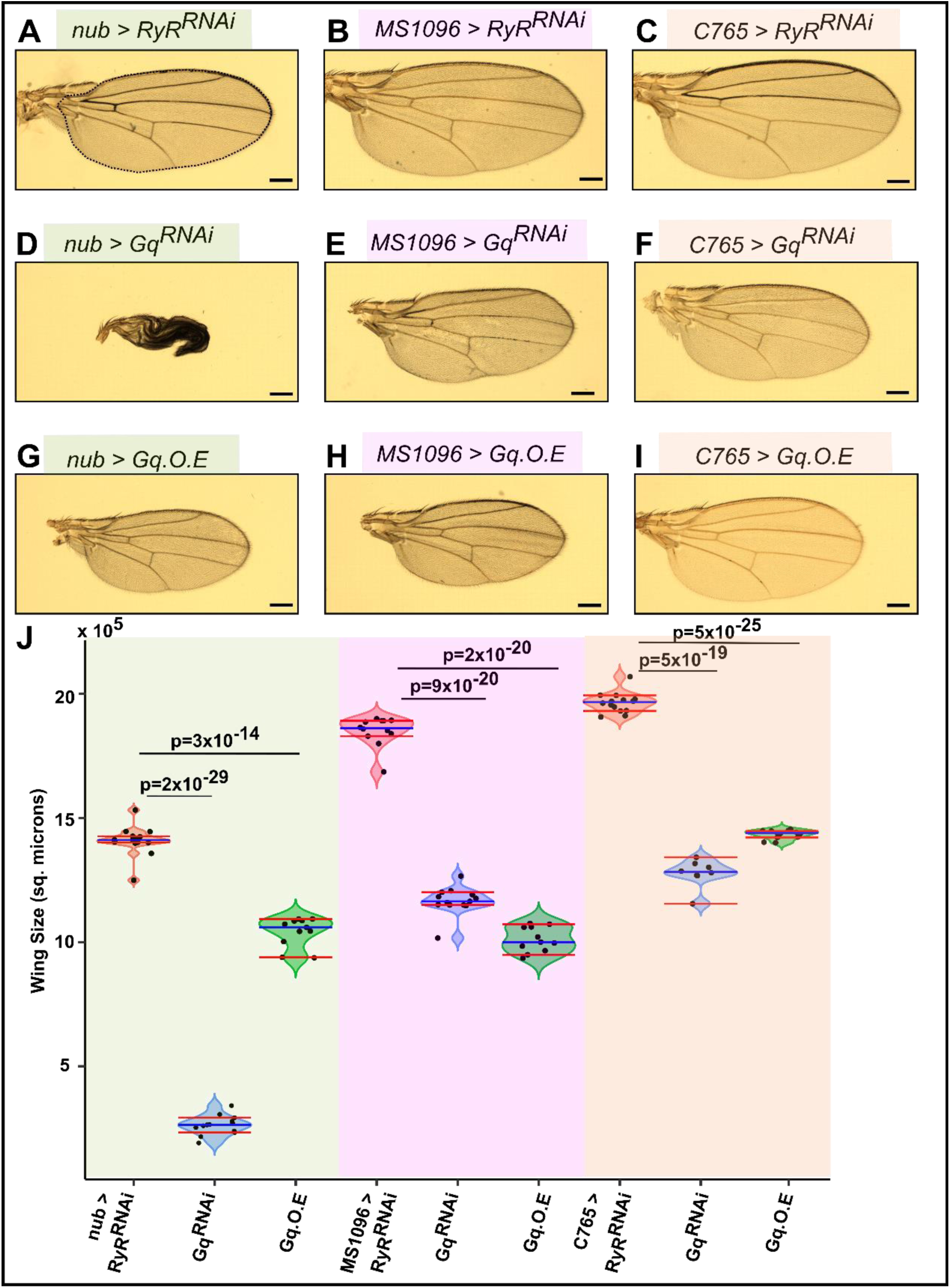
Perturbing *Gαq* expression in the *Drosophila* wing disc decreases overall adult wing size and can prevent pupal wing expansion. **A-I)** Micrographs of adult wings expressing *RyR^RNAi^* (control), wild type *Gαq*, or *Gαq*^RNAi^ under several GAL4 drivers, including nubbin, MS1096, and C765. **D-F)** Knockdown of *Gαq* leads to a reduction in wing size for multiple Gal4 drivers. RNAi-mediated inhibition of *Gαq* driven by the *nubbin-Gal4* driver led to severe pupal wing expansion phenotypes. **G-I)** *Gαq* overexpression leads to decreased wing size for multiple drivers. **J)** Graphs comparing the wing area for wings from the crosses indicated in (A-I). The blue line represents the median, and the red lines indicate the 95% confidence interval of the median. N > 10 for all samples shown except for *nub> Gαq^RNAi^* (n=8). The scale bar represents 100 microns. The blue line represents the median, and the red lines indicate the 95% confidence interval of the median. Student two-tailed t-test was used to determine the statistical significance.

### Inhibition of Ca^2+^ signaling enhances *Gαq* overexpression phenotypes

Next, we investigated the role of Gαq mediated calcium signaling towards the growth inhibition caused by *Gαq* overexpression. Our previous findings demonstrated that the downregulation of Ca^2+^ signaling components using the MS1096-Gal4 driver resulted in a size reduction of the adult wing[67]. To confirm this, we downregulated the Ca^2+^ signaling components, including *small wing* (*sI: a homolog of PLCγ)*[72] and *IP_3_R*, using a C765-Gal4 driver (**Fig. 3A, E, G, I**). The release of Ca^2+^ from ER to the cytoplasm is dependent on the binding of inositol trisphosphate (IP_3_) to the inositol trisphosphate receptor (IP_3_R). Further, IP_3_ is produced by PLC through the hydrolysis of Phosphatidylinositol 4,5-bisphosphate (PIP2). Thus, a major mode of Ca^2+^ signaling relies on IP_3_ and PLC activity. Consistent with our earlier findings, we observed a reduction in the wing size for the knockdown of Ca^2+^ signaling components. Additionally, the knockdown of PLCβ homolog Phospholipase C at 21C *(Plc21C)* reduced the final wing size (**Fig. 3C, I**). These results imply that Gαq-mediated Ca^2+^ signaling promotes growth during organ development.

**Figure 3:**
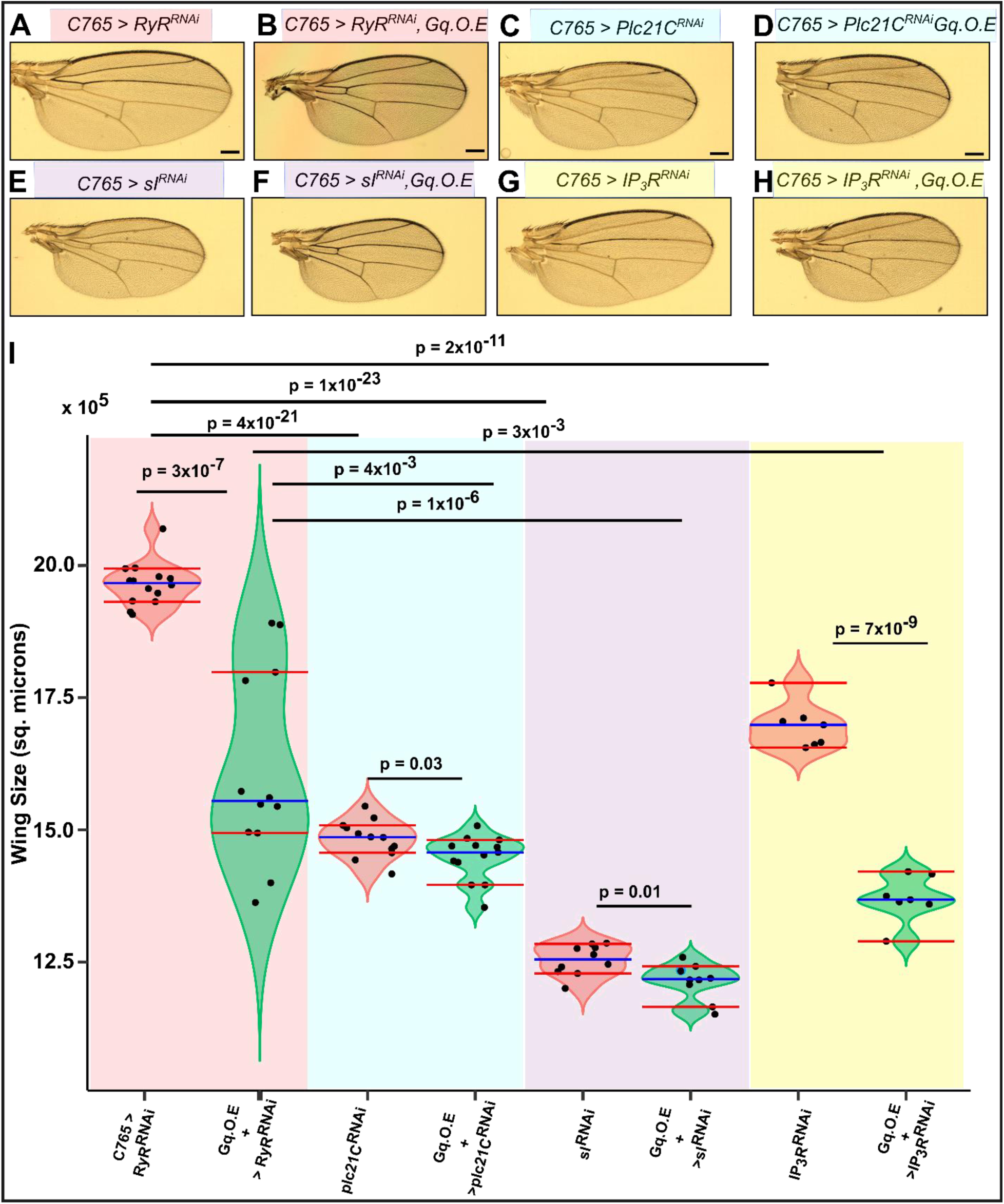
*Gαq* overexpression, together with inhibition of Ca^2+^ signaling components, results in further reduction of wing size. **A-H**) Micrographs of adult wing images. Reduction in core Ca^2+^ signaling components such as *plc21c*, *sI*, and *itpr* (IP_3_R) through RNAi-mediated knockdown in the wing disc results in a reduction of the size of the wings. Overexpressing wildtype *Gαq*, in combination with the knockdown of Ca^2+^ signaling components, further decreases the size of the wings. **I**) Quantification of the adult wing blade areas for the crosses listed in (**A-H**). The blue line represents the median, and the red lines indicate the 95% confidence interval of the median. Scale bars are 100 µm. Student t-test was used to determine the statistical significance.

In contrast, *Gαq* overexpression, which increases Ca^2+^ activity in the wing disc, reduces the final wing size. As a next step, we investigated whether regulators of Ca^2+^ activity contribute to the reduction in wing size. To answer this, we simultaneously overexpressed wildtype *Gαq* and downregulated Ca^2+^ signaling components, such as *PLCs* and *IP_3_R,* using RNAi. We observed that the *Gαq* overexpression, along with the knockdown of Ca^2+^ signaling components, enhanced the reduction in wing size (**Fig. 3A-I**). This double perturbation analysis suggests that Gαq- mediated Ca^2+^ signaling promotes growth, whereas growth inhibition due to *Gαq* overexpression occurs through a mechanism independent of Ca^2+^ signaling. Altogether, our results imply that Gαq plays a dual role, where it promotes growth through a Ca^2+^-dependent and inhibits growth through a Ca^2+^-independent mechanism.

### Perturbation of Gαq levels resulted in increased cell density and decreased cell number

To examine whether Gαq mediates growth by regulating cell proliferation, we performed immunohistochemistry to measure Phospho-Histone H3 (PH3) levels in the wing discs. We quantified the PH3 levels in the wing disc by calculating the proliferation density. We measured the proliferation density by dividing the total area of the nuclei positive for PH3 by the pouch area (**Fig. 4**). Our analysis revealed that the cell proliferation density (PH3 per unit area of the wing disc) for both *Gαq* perturbations (overexpression or RNAi mediated inhibition) was higher than those of the control condition. Moreover, we identified that the wing discs were smaller in both *Gαq* perturbations (overexpression and downregulation) compared to the control condition (**Fig. 4D**). (**Fig. 4E**). Next, we analyzed the trichome number and trichome density, as the trichome number indirectly represents the total number of cells in the wing. Our analysis revealed that the trichome number decreased significantly with the knockdown of *Gαq* (**Fig. 4G**), whereas *Gαq* overexpression showed no change in the trichome number. However, the trichome density was significantly higher for both overexpression and knockdown conditions (**Fig. 4I**). An increase in the trichome density (**Fig. 4I**) indicates that the average cell area is smaller for both *Gαq* perturbations. Cumulatively, these results indicate that *Gαq* overexpression reduces cell size, whereas *Gαq* knockdown reduces cell number and size.

**Figure 4:**
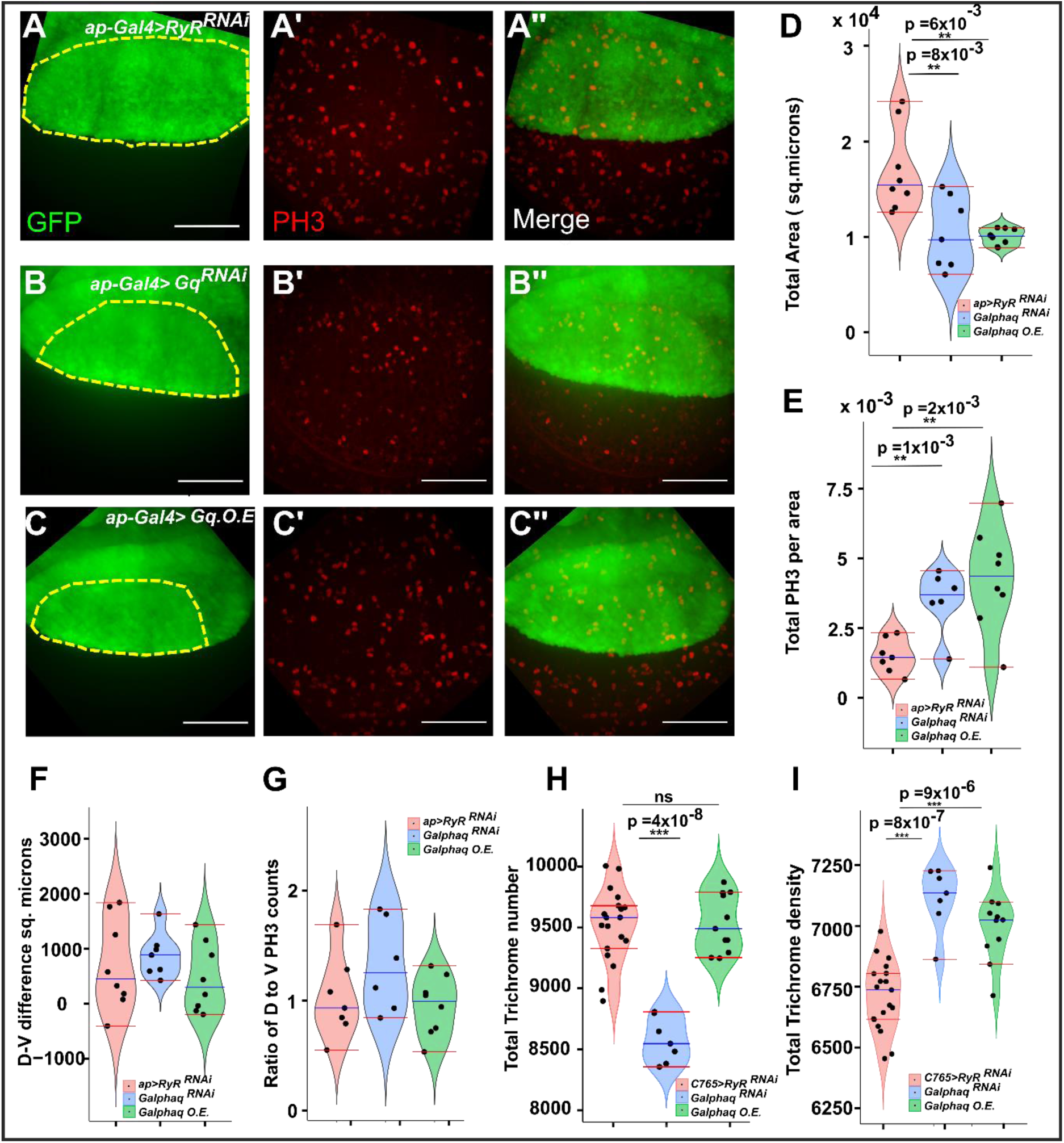
Gαq homeostasis modulates the proliferation of wing disc and affects the overall cell number in the wing. **A**, **B**, **C**) Representative images of wing discs stained with PH3 expressing RyR^RNAi^, *Gαq*^WT,^ and *Gαq* ^RNA^_i_ under the control of apterous. PH3 density was higher in the discs with *Gαq* overexpression and RNAi conditions when compared to the control. **D**) Violin plots comparing the total area of the wing pouch for *Gαq* perturbations with control. The pouch area was significantly reduced in the overexpression and RNAi conditions as compared to the control condition. **E**) Violin plot showing the density of PH3 in the wing pouch for the control and *Gαq* perturbations. PH3 density was found to be higher in both *Gαq* overexpression and RNAi conditions. **F**) The violin plots illustrate the difference between the area of the dorsal and ventral regions of the wing discs under control and *Gαq* perturbations. **G**) There exists no difference in the number of PH3 cells in the dorsal and ventral regions for all the genetic perturbations. (n>5 for all conditions) **H**) Violin plot comparing trichome number between wings with *Gαq* overexpression, RNAi, and control conditions. Trichome number is significantly lower for wings with *Gαq* knockdown, whereas overexpression does not show any difference. **I**) Graphs comparing the total trichome densities for *Gαq* overexpression, *Gαq*_RNAi_, and *RyR_RNAi_*. Trichome density is significantly higher for both overexpression and RNAi conditions indicating that Gαq homeostasis affects the cell number during the growth of the wing disc.

Next, we extended the trichome analysis for individual intervein regions to investigate the effects of *Gαq* perturbations on intervein regions. Our intervein analysis showed that the trichome number decreases in all the intervein regions for the *Gαq* knockdown. In contrast, the trichome number increased for *Gαq* overexpression in the intervein regions 3, 4, and 5 (as defined in **Additional File 1, Fig. S1A**). Hedgehog (Hh) signaling has been associated with regulating the growth and patterning of the L3-L4 region of the wing, which corresponds to the intervein region 4[73]. Further, our trichome density analysis shows a different trend in the intervein regions 3 and 5. We observed that the trichome density did not show any significant difference in intervein region 5 for either inhibition of *Gαq* or overexpression of *Gαq* (**Additional File 1, Fig.S1B**). Intervein regions 3 and 5 are associated with high levels of Hedgehog (Hh) signaling and Decapentaplegic (DPP) signaling during larval development, respectively[74] (**Additional File 1, Fig.S1B**). Together, these results confirm that perturbation to Gαq levels impacts cell proliferation and growth, thus affecting the final cell number and cell size. However, this effect is also dependent on the background signaling in different wing regions.

### Caspase-mediated apoptotic response in *Gαq* perturbations does not influence wing size

To determine whether apoptosis is stimulated by perturbing levels of *Gαq*, we performed immunostaining for death caspase-1 (*DCP-1*), a marker of programmed cell death (**Fig. 5A**). As caspases play an essential role in apoptosis[75], we quantified the area of *Dcp-1* expressing cells in both *Gαq* overexpression and knockdown conditions. Surprisingly, we discovered that either increasing or decreasing the levels of Gαq slightly decreased the Dcp-1-induced apoptotic cells in the wing disc. However, this effect was not statistically significant, given the relative background levels of Dcp-1 staining (**Fig. 5A, F**). To test whether Gαq-mediated apoptotic response determines the reduced wing size, we co-expressed both wildtype *Gαq* and baculovirus *p35* in the larval wing disc using the C765-Gal4 driver and assayed for the final adult wing phenotype. Baculovirus *p35* inhibits cell death in developing eyes and fly embryos[76] and inhibits caspases[75]. Co-expression of *Gαq*, along with p35, could not rescue the wing blade reduction. Interestingly, we observed a further reduction in the wing blade area when both *p35* and *Gαq* were expressed (*C765 > Gαq;+ > p35*) in the wing disc (**Fig. 5B-E**). In summary, inhibition of apoptosis by ectopic expression of apoptosis inhibitor *p35* did not rescue wing size reduction due to overexpression of *Gαq*. Our results thus indicate that caspase-mediated apoptotic response is not a major determinant in reducing the size of wings when *Gαq* levels are perturbed.

**Figure 5:**
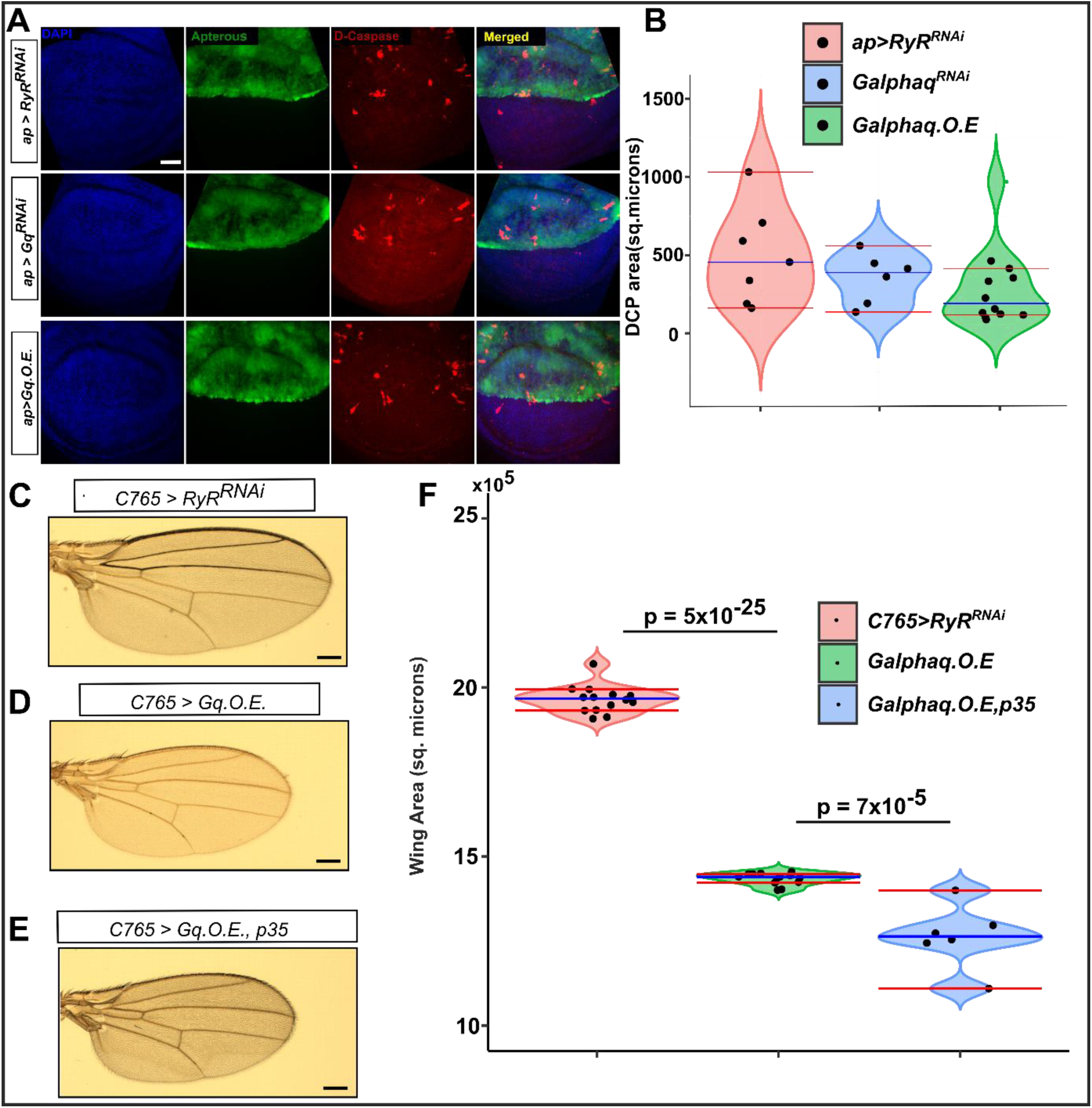
Reduction in adult wing size for both *Gαq* overexpression and downregulation is not mediated by apoptosis. **A)** Representative images of the third instar wing disc from the corresponding cross stained for Dcp-1 to show apoptotic cells. Wing discs were also stained with nuclear dye DAPI, and the region of GAL4 expression, which drives the *Gαq* perturbation, is indicated using GFP. All wing discs are positioned anterior to the right and posterior to the left. Sample sizes are n = 3 to 8 for each condition. **B**) Violin pot comparing the area of Dcp-1 stain for all three genotypes. **C-E**) Micrographs of adult wings. Genotypes of individual wings are given on the top left. Ectopic expression of apoptosis inhibitor p35 did not rescue the size reduction defect caused by overexpression of *Gαq*. **F**) Quantification of the adult wing blade area from the indicated genotypes. Student t-test was performed, and the p-values are indicated on the plot. The blue line represents the median, and the red lines indicate the 95% confidence interval of the median. Scale bars represent 100 µm.

### *Gαq* overexpression stimulates stress response genes

To examine the transcriptional profile activated by *Gαq* overexpression, we performed RNA-seq on control discs expressing a control construct (*RyR^RNAi,^* which has no known effect on wings) and discs overexpressing *Gαq* (under the control of the *C765-Gal4* driver). Compared to the control discs, overexpression of *Gαq* with the C765-Gal4 driver resulted in 3613 differentially expressed genes with an adjusted p-value (p-value adjusted using Benjamini-Hochberg) of less than 0.05 (**Additional File 1, Fig. S2A**). Of these genes, 2009 had a fold change greater than 1.5, with 878 genes upregulated and 1131 genes downregulated. To ascertain the biological processes affected by the differentially expressed genes, we performed a Gene Ontology (GO) enrichment analysis of the genes having an absolute log fold change greater than 0.6 (abs (log2FC ≥0.6)). Based on our GO analysis, we found enrichment in biological processes related to DNA replication, cuticle development, molting cycle, myofibril assembly, and protein refolding (**Additional File 1, Fig. S2B**). To investigate the biological processes that were activated or suppressed, we examined the significantly upregulated and downregulated genes separately.

First, we performed a GO enrichment analysis of the upregulated genes having a fold change greater than 1.5 (log2FC≥0.6) (**Additional File 1, Fig. S3A**). According to our GO analysis, we identified that the upregulated genes are involved in biological processes related to DNA replication, ribosome biogenesis, chromosome segregation, mitotic cell cycle process, and cuticle development. Moreover, we examined the relationships between the top enriched GO terms (**Additional File 1, Fig. S3B**) and identified that the top GO terms cluster under DNA replication, ribosome biogenesis, and cuticle development (**Additional File 1, Fig. S3B**). Additionally, we examined the genes associated with the enriched biological processes and their respective fold change values. We observed that the majority of the upregulated genes were related to the DNA replication and the mitotic cell cycle process, consistent with the possibility that the wing discs were physiologically younger and were proliferating (**Additional File 1, Fig. S3C**). Upon removing the redundant GO terms, we found that the upregulated genes are also involved in biological processes related to muscle cell development, non-membrane-bound organelle assembly, actomyosin structure organization, and protein refolding (**Additional File 1, Fig. S4**). Moreover, we found upregulation in heat shock proteins (*Hsp 70Ba*), which is related to the GO terms such as protein refolding (**Additional File 1, Fig. S4**). As well we observed an upregulation in myosin heavy chain (*Mhc*) and myosin light chain (*Mlc2*), which are involved in the GO terms such as muscle cell development and actomyosin structure organization (**Additional File 1, Fig. S4**).

Our analysis further revealed the downregulation of Serine Protease Inhibitors (Serpin 100A, Spn 100A) (**Additional file 2**) as well as an increase in serine proteases, including multiple Jonah genes (**Additional File 1, Fig. S5**). Serine proteases are linked to the GO terms such as serine hydrolase, peptidase activity, hydrolase activity, and endopeptidase activity (**Additional File 1, Fig. S5**). Serine proteases initiate proteolysis, thus eliciting various immune responses to infection[77]. Moreover, Jonah genes are expressed during larval development and exclusively in the midgut[78]. In addition, we observed that the upregulated heat shock proteins were also associated with the hydrolase activity (**Additional File 1, Fig. S5**). Activation of heat shock proteins is a defense mechanism employed by cells in response to stress caused by the accumulation of abnormal or denatured proteins[79], [80] (proteins having defects in folding). Our GO analysis supports this result as we observe an enrichment in biological processes related to protein refolding and folding (**Additional File 1, Fig. S4**). Upregulation of heat shock proteins, particularly *Hsp70*, implies the possibility of stress accumulation in the cells. Further, induction of *Hsp70* proteins is known to reduce the growth rate in *Drosophila* cells[81, p. 70], which is potentially consistent with reduced cell size phenotypes and associated reduction in overall wing growth.

Additionally, our RNA seq analysis revealed upregulation of genes associated with chitin-based cuticle development, cuticle development, structural constituent of the cuticle, encapsulating structure, and extracellular matrix (**Additional File 1, Fig. S3-S5**). An important consideration is that the amount of cuticle proteins produced by a larva varies along with its developmental stage. Furthermore, cuticle protein synthesis depends on the 20-hydroxyecdysone (20E) levels and occurs during periods of decline in ecdysteroid levels[82]. Moreover, our analysis revealed an increase in Dilp8 RNA levels, which also corresponds to the GO term external encapsulating structure (**Additional File 1, Fig. S5**). Dilp8 is an insulin-like peptide regulating the timing of growth and, thus, the larval-to-pupal transition[83], [84]. In summary, our analysis identified that *Gαq* overexpression increased the activity of serine proteases and heat shock proteins and affected cuticle protein synthesis. Based on our GO enrichment analysis, we conclude that the *Gαq* overexpression upregulates genes involved in DNA replication, protein folding, and cuticle protein synthesis.

As a next step, we extended our enrichment analysis to the downregulated genes (log2FC ≤ -0.6) (**Additional File 1, Fig. S6**). Enrichment analysis revealed that the significantly downregulated genes mainly correspond to biological processes such as the molting cycle, axon development, larval development, cell adhesion, and cell morphogenesis (**Additional File 1, Fig. S6A**). Upon examining the similarity between the top GO terms, we identified that locomotion, axon guidance, and cell morphogenesis were closely related (**Additional File 1, Fig. S6B**). Additionally, we examined the downregulated genes that are related to the enriched GO terms. We observed that most of the genes downregulated were involved in locomotion, taxis, neuron projection development, and biological adhesion (**Additional File 1, Fig. S6C**). We identified that the cell death genes such as *head involution defective*(*hid*) and *Dronc* were downregulated and mapped to the GO term neuron projection development. In addition, we identified downregulation in cuticular proteins (*Cuticular protein 31A*, *Cuticular protein 72Ea*, etc.) and genes related to the molting cycle (**Additional File 1, Fig. S6C**, **Fig. S7**). Moreover, our analysis revealed a strong downregulation in 20-hydroxyecdysone (20E) target genes, which include *Hormone receptor 4 (Hr4)*, *Blimp-1, and Ecdysone-induced gene 71Ee (Eig71Ee).* (**Fig. 6B&C, Additional File 1, Fig. S7C, Fig. S8**). Since ecdysone plays a pivotal role in the synthesis of cuticle proteins, we interpret that the low levels of ecdysone signaling might affect the deposition of cuticle proteins and their timing of deposition. Collectively, these results demonstrate that *Gαq* overexpression inhibits ecdysone signaling and, as a result, affects cuticle protein synthesis.

**Figure 6:**
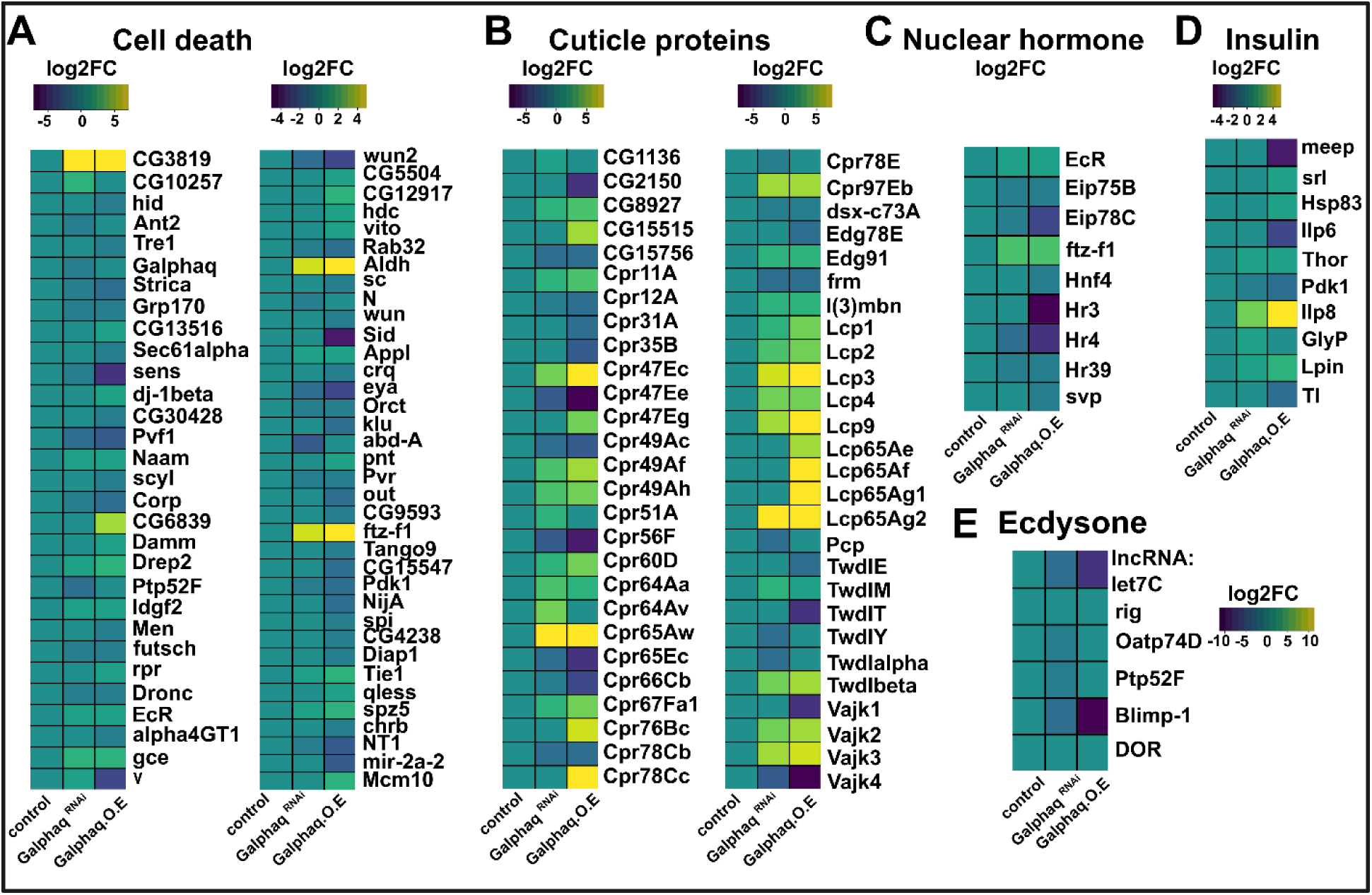
Disruption of Gαq homeostasis delays development through transcriptional changes in multiple modules. **A**) Transcriptomic analysis of genes related to programmed cell death for *Gαq* RNAi and *Gαq* overexpression. The apoptotic gene (*ftz-f1*) and Mitochondrial apoptotic genes (*CG3819* and *Aldh*) were upregulated, and negative regulators of apoptosis (*sens*, *eya*, *Pvf1*, *futsch*) were downregulated in both conditions. Also, cell survival genes such as *spitz*, an EGFR ligand, were downregulated. **B**) Strong dysregulation of cuticle proteins is observed in both *Gαq* overexpression and *Gαq* RNAi conditions. **C**) Ecdysone genes (*Blimp-1*, *IncRNA: let7c*) and nuclear hormone receptors (*Eip78c*, *Eip75b*, *Hr3*, *Hr4*) were downregulated. *Ftz-f1* and *ECR*, which are expressed in early larval stages, were upregulated. **D**) Heat map plot showing gene expression changes of the Insulin signaling components. *Pdk1* was downregulated in both *Gαq* O.E. and RNAi perturbations. Relaxin-like *dilp8* was significantly upregulated in both *Gαq* overexpression and RNAi conditions.

### *Gαq* downregulation inhibits ecdysone-induced genes

To determine the transcriptomic responses affected by the *Gαq* knockdown, we performed RNA sequencing on the wing discs expressing *Gαq*^RNAi^ and the control discs expressing *RyR^RNAi^*. Compared to the control, 2667 genes were differentially expressed with an adjusted p-value less than 0.05. Of these genes, 1173 had absolute fold change greater than 1.5, of which 555 genes were upregulated, and 618 genes were downregulated (**Additional File 1, Fig. S9A**). To determine the biological processes enriched among the differentially expressed genes, we performed a gene ontology enrichment analysis of the genes that have a significant fold change (abs (log FC) ≥ 0.6). GO classification analysis revealed that the differentially expressed transcripts were dominant in biological processes related to transmembrane transport, cuticle development, molting cycle, and chitin-based cuticle development (**Additional File 1, Fig. S9B**). Surprisingly, the enriched GO terms for the RNAi were mostly like those observed in the *Gαq* overexpression condition. Both overexpression and downregulation of *Gαq* resulted in the enrichment of biological processes related to the cuticle proteins, axon development, and molting cycle (**Fig. 6B, Additional File 1, Fig. S9B**).

Next, we performed the Gene Ontology enrichment analysis of the significantly upregulated genes with a fold change greater than 1.5 (log FC ≥ 0.6). Our GO analysis of upregulated genes showed enrichment in biological processes related to cuticle development, mitotic spindle organization, and DNA replication (**Additional File 1, Fig. S10A**). After examining the upregulated genes relevant to the enriched GO terms, we identified that the upregulated genes were mainly involved in DNA replication and cuticle development. For instance, genes such as *Orc2* (*Origin recognition complex subunit 2*), *Minichromosome maintenance 2* (*Mcm2*), and *Proliferating cell nuclear antigen* (*PCNA*) were upregulated and mapped to the GO terms such as DNA replication and DNA dependent DNA replication (**Additional File 1, Fig. S10C**). In addition, we identified upregulation in cuticle proteins such as *Cuticular protein 65Aw (Cpr65Aw*, *Cpr60D, etc.),* which are associated with the GO term cuticle development (**Additional File 1, Fig. S10C**). In addition, our analysis revealed an upregulation in serine proteases which map to the hydrolase activity and peptidase activity (**Additional File 1, Fig. S11**). Overall, our analysis of upregulated genes revealed that *Gαq* knockdown disrupted cuticle protein synthesis.

We then conducted a GO enrichment analysis of the downregulated genes to determine the biological processes associated with the downregulated genes. We identified that the downregulated genes enrich the GO terms associated with transmembrane transport, molting cycle, cell recognition, axon guidance, neuron projection guidance, and mating behavior (**Additional File 1, Fig. S12A**). After examining the gene expression relevant to the enriched GO terms, we identified that most of the downregulated genes were associated with cuticle development, molting cycle, transmembrane transport, taxis, and axon development. Interestingly, our analysis revealed downregulation in genes involved in ecdysone response regulation, including *ecdysone-induced protein 75*B (*Eig75*), which mapped to the GO terms cuticle development and molting cycle[85]. As our study revealed downregulation in genes mediating ecdysone response, we examined the gene expression related to ecdysone signaling. Interestingly, we found that the downregulated ecdysone response genes predominantly belong to the *ecdysone-induced genes 71E* family (*Eig71E*), which also are associated with the defense and immune response (**Fig.6D, Additional File 1, Fig. S13**).

*Eig71E* consists of five late-puff gene pairs that are transcribed divergently[86]. Moreover, the *Eig71E* family encodes small peptides that have biochemical characteristics similar to vertebrate defensins and are thus assumed to provide an antimicrobial defense during metamorphosis[86]. These genes are induced by the expression of early puff genes, which are directly regulated by the steroid hormone 20-hydroxyecdysone. As the *Eig71* family belongs to late puff genes, their expression would be moderate in L3 and increase in late L3. Our data show a strong downregulation of *Eig71E* genes for *Gαq*^RNAi^ compared to the control (**Additional File 1, Figure S13**). Overall, this result suggests that the wing discs expressing *Gαq^RNAi^* were physiologically less mature when compared to the control discs. Hence, we observed a downregulation in the late puff genes (**Fig. 6F**), as their expression is restricted to late larval stages. As the expression of these *Eig71E* genes is induced by ecdysone, we hypothesize that downregulation in ecdysone signaling for the discs expressing *Gαq^RNAi^* (**Fig. 6C**). Compatible with this hypothesis, we observed a downregulation in ecdysone genes such as *Hr4* and *Blimp-1* (**Fig. 6C**). The results of our study also indicated a downregulation of *Organic anion transporting polypeptide 74D* (*Oatp74D*), a solute carrier transporter that is responsible for transporting ecdysteroids into cells[87] (**Additional File 1, Fig. S12**). We further identified that the genes related to cell death were downregulated (**Additional File 1, Fig. S13**). Most genes related to cell death have a significant fold change for both overexpression and RNAi conditions (**Fig. 6A**). In summary, our study showed that the downregulation of *Gαq* in the wing disc inhibits ecdysone pathway genes **(Additional File 1, Fig. S13**).

### Enrichment analysis of major signaling pathways

We next examined how *Gαq* perturbations affect the core signaling pathways that play imperative roles in growth and development. To achieve this, we performed an enrichment analysis to investigate the core signaling pathways that were significantly represented among the differentially expressed genes with statistically significant fold changes[88]. Our analysis identified that the Toll signaling and nuclear hormone receptor pathway were significantly enriched among the differentially expressed genes in the *Gαq* overexpression condition (**Fig.7A, B**). In the case of *Gαq^RNAi^,* the nuclear hormone receptor pathway was enriched among the differentially expressed genes. In addition, we analyzed upregulated and downregulated genes separately to identify the enrichment of core pathways among them. We found that the JAK/STAT pathway was significantly enriched among the upregulated genes for the *Gαq* overexpression, and no pathway was enriched among the upregulated genes for *Gαq^RNAi^*. Next, our analysis revealed that the nuclear hormone and the Toll signaling pathway were enriched among the downregulated genes for *Gαq* overexpression. For *Gαq^RNAi^*, the nuclear hormone receptor pathway was significantly enriched among the downregulated genes. In summary, our enrichment analysis revealed that the nuclear hormone receptor, JNK/STAT, and Toll pathways were significantly enriched for *Gαq* OE, whereas the nuclear hormone receptor pathway was significantly affected for *Gαq^RNAi^* (**Fig 7A, B**).

**Figure 7:**
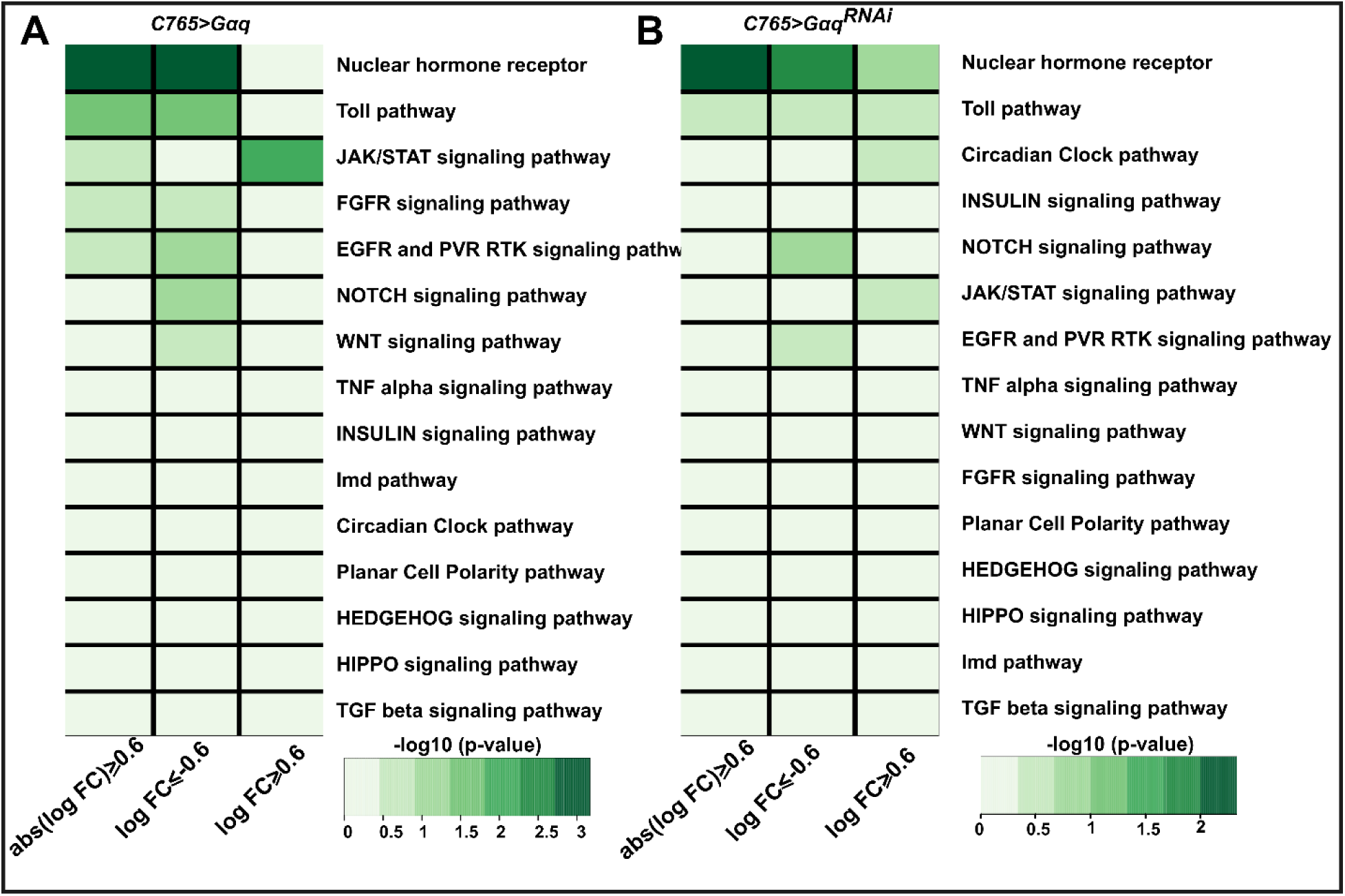
Enrichment analysis of core signaling pathways among the differentially expressed genes. **A**) Overexpression *Gαq* resulted in the enrichment of the Toll pathway and Nuclear hormone receptor among the differentially expressed genes with a significant fold change (abs (log2FC) ≥ 0.6). Toll pathway and nuclear hormone receptor pathway were enriched among the genes that were significantly downregulated (log2FC ≤ -0.6). JAK/STAT signaling was enriched among the significantly upregulated genes (log2FC ≥ 0.6). **B**) For the *Gαq_RNAi_* perturbation, the nuclear hormone receptor pathway was enriched among the differentially expressed genes with a significant fold change (abs(log2FC) ≥ 0.6). Nuclear hormone receptor pathway was enriched among the significantly downregulated genes. No core signaling pathways were enriched among the significantly upregulated genes (log2FC ≤ -0.6).

Following this, we examined gene expression changes associated with specific components of the core signaling pathways that were affected. As a first step, we examined changes in the gene expression of Toll signaling components. Toll signaling regulates cellular immune response and mediates cell elimination by regulating growth and apoptosis[89], [90]. Our analysis showed a significant upregulation in the *Toll-9* for both *Gαq* perturbations (**Additional File 2**). Additionally, we observed an approximate two-fold upregulation in *spatzle-4* (*spz4*) for *Gαq^RNAi^* and *spatzle-5* (*spz5*) for *Gαq* OE perturbations. However, we observed a downregulation in *spatzle* (*spz*) for *Gαq* OE condition (**Additional File 1, Fig. S14**). Cleaved Spatzle is known to activate the Toll pathway in unfit cells during cell competition, thereby activating cell death[89], [91]. Upregulation of *spatzle-4* and *spatzle-5* in our analysis likely resulted from the upregulation of serine proteases. Serine proteases induce proteolytic cleavage of Spatzle during an immune response resulting in the Toll activation[92]–[94]. Cleaved Spatzle binds to the Toll receptor and initiates the nuclear localization of *Dorsal-related immunity factor* (*Dif*), resulting in the transcription of several immunity genes. Our RNA seq results showed an increase in fold change values of *Dif* for both perturbations (**Additional File 1, Fig. S14, Fig. S15**)[95]. Of note, studies have shown that activation of Toll signaling in fat body cells inhibits insulin signaling in a non-autonomous manner, thereby resulting in reduced body growth [96]–[98]. Our analysis revealed a strong downregulation in *Phosphoinositide-dependent kinase 1* (*Pdk1*) for both *Gαq* overexpression and downregulation (**Fig. 6D, Additional File 1, Fig. S14, Fig. S15**). *Pdk1* acts upstream of *Akt* phosphorylation, and Toll signaling inhibits *Akt* phosphorylation in fat body cells[96], [98]. Normally, microbial peptides and unmethylated CpG DNA sequences activate the Toll-9 receptor. Our data shows upregulation in heat shock proteins for both *Gαq* perturbations, which can induce the activation of Toll receptor*s*. As *Gαq* perturbations affected the expression of multiple components of Toll signaling, we propose that Gαq might play roles in Toll signaling, thus possibly affecting growth and apoptosis.

As Toll signaling activates an immune response, we asked whether immune-related genes were affected. We examined the fold change values of the immune signaling components in both *Gαq* overexpression and *Gαq*^RNAi^ perturbations (**Additional File 1, Fig. S14, Fig. S15**). We observed a downregulation in the immune response genes for both perturbations, except for the genes including *Immune induced molecule 33* (*Im33*), *Lysozyme S* (*LysS*), and *Lysozyme B* (*LysB*). *IM33* is strongly upregulated for both perturbations and could act downstream of the Toll pathway.

Since upregulated genes in *Gαq* OE affected JAK/STAT signaling, we examined the expression change in specific components of JAK/STAT signaling. We observed that *unpaired 3* (*upd3*) was upregulated with *Gαq* overexpression with a three-fold change (**Additional File 1, Fig. S14**). *Unpaired3* (*upd3*, interleukin-6-like cytokine) is implicated in immunity, repair, gene expression, and development[99]–[101] through its activation of the JAK/STAT signaling pathway. In addition, *chinmo* (*chronologically incorrect morphogenesis),* a downstream effector of JAK/STAT signaling[102], is significantly upregulated in both *Gαq* ^RNAi^ (∼12 fold) and *Gαq* OE (∼46 fold) (**Additional File 2**). Further, we identified that both perturbations increased the expression of JAK/STAT target *dilp-8* (**Fig. 6D**). The release of Dilp-8 is triggered by the activation of JAK/STAT signaling, which leads to ecdysone inhibition, thereby slowing the growth[103]. Furthermore, Dilp8 governs larval-to-pupal transition and developmental timing by regulating ecdysone signaling[84]. In summary, our study identified that *Gαq* perturbations change the expression of multiple JAK/STAT signaling components, which may contribute to either growth or apoptosis[104], [105].

As *Gαq* perturbations affected the nuclear hormone receptor pathway, we examined the fold change values of nuclear hormone receptors. Nuclear receptors bind directly to DNA and modulate gene transcription. Our data showed that the majority of the nuclear hormone receptor genes were downregulated (**Fig. 6C**), except for *fushi tarazu transcription factor 1* (*ftz-f1*) and *Ecdysone receptor* (*EcR)* (**Additional File 1, Fig. S14, Fig. S15**). Also, we observed a downregulation in ecdysone-induced genes for both perturbations. *Ftz-f1* is a nuclear receptor that transcriptionally regulates the synthesis of cuticle proteins and controls metamorophosis[82], [106]. Additionally, *EcR* is a nuclear receptor that mediates the tissue’s response to ecdysteroids, which is integral to larval molting and metamorphosis[107]–[109]. *Ftz-f1* is a mid-prepupal puff gene, which also is expressed in the early second instar stage of larval development[106], [110]. In addition, *EcR* is expressed during the second half of the second instar stage of larval development[108], [110]. Thus, upregulation in *ftz-f1* and *EcR* likely resulted from the delay in larval development caused by *Gαq* perturbations. Taken together, our study indicates that perturbing Gαq levels disrupt larval development by affecting ecdysone signaling and nuclear hormone expression.

As noted above, *dilp8* expression levels were significantly upregulated. So, we asked whether this increase in *dilp8* impacts the larval to pupal transition time when *Gαq* expression levels are perturbed[103]. To determine this, we performed a developmental timing assay, wherein we recorded the larval to pupal transition time for both control and *Gαq* perturbations. Consistent with the RNA seq finding, we observed that the larvae with *Gαq* overexpression in the wing disc cells pupariated 24 h later than the wildtype larvae (**Fig. 8A**). Similarly, we observed a delay in larval to pupal transition by 12 h for *Gαq* knockdown (**Fig. 8A**). Overall, these results imply that the disruption of Gαq levels inhibits the ecdysone signaling, therefore, delaying the development to compensate for the reduced growth. One likely interpretation of these results is that *Gαq* overexpression in the wing disc elicits immune and stress response through activation of the JAK/STAT pathway. This might activate the heat shock proteins that decrease the growth rate of the wing. JAK/STAT stimulates *dilp-8* production, which signals the brain to inhibit ecdysone synthesis, thereby slowing down development. For the *Gαq*^RNAi^ data, we interpret that decrease in Ca^2+^ activity caused by *Gαq* downregulation leads to a reduction in growth. Calcium is an important second messenger that regulates multiple cellular responses, including growth and apoptosis[26], [67], [111]. Hence, we propose that the downregulation of *Gαq* affects growth through reduced Ca^2+^, thus negatively affecting ecdysone signaling. Strikingly, we observed Ip_3_R inhibition, along with *Gαq* overexpression, reduced the developmental delay (**Fig 8A**). As a result, this finding supports the hypothesis that increased calcium levels are responsible for increasing the growth time that occurs through the delay in larval to pupal transition when *Gαq* is overexpressed.

**Figure 8:**
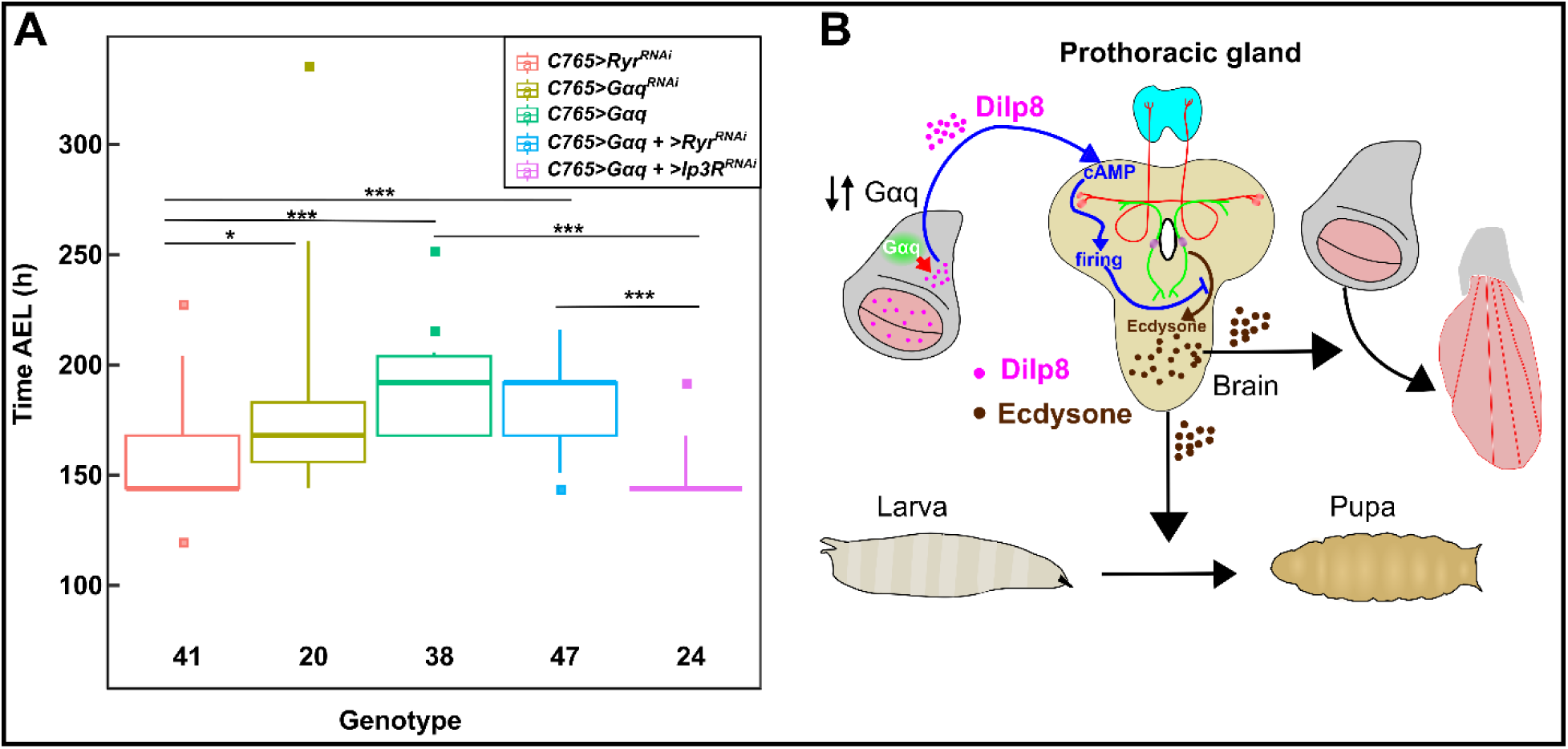
Disruption of Gαq homeostasis delays larval to pupal transition. **A**) A box plot illustrating the larval to pupal transition time for the genetic perturbations shown in the panel. Whiskers represent 5% and 95% percentiles. Square points represent the outliers. A Kruskhal- Wallis test revealed significant differences in pupariation times between the *Gαq* perturbations (the degrees of freedom is 4, the H is 62, and the p-value of 9.16 x 10_-13_) (Degrees of freedom represents the number of groups subtracted with one. The test statistic H is compared with the critical value Chi-square for the given degrees of freedom). Conover post hoc test was used to perform multiple pairwise tests to compare the pupariation times between different genotypes. Asterisks indicate genotypes with statistically significant differences in pupariation times. Overexpression of *Gαq* delays larval to pupal transition by ∼24 h, whereas *Gαq_RNAi_* delays transition time by ∼12 h. Knockdown of Ip_3_R along with the *Gαq* overexpression rescues the developmental delay observed with *Gαq* overexpression. **B**) Proposed mechanism of developmental delay: Dysregulation of Gαq homeostasis in the wing disc upregulates *dilp-8* expression, ultimately resulting in decreased 20E synthesis from the brain. This generalization is consistent with RNA seq analysis from this work, where overexpression and downregulation of *Gαq* in the wing disc result in the downregulation of 20E target genes, respectively. Figure inspired from [157].

## Discussion

Extracellular signals regulate organ development by embedding encoded information into the spatiotemporal dynamics of second messenger concentrations. G proteins play a vital role in mediating these external signals from GPCRs to downstream effectors by regulating key second messengers, including Ca^2+^, cAMP, and IP_3_, among others. Recent work has demonstrated that the spatiotemporal dynamics of Ca^2+^ signaling correlate with changes in the final wing size in diverse ways, ranging from single-cell repetitive spiking to global traveling multicellular waves[67]. In particular, the G-protein Gαq stimulates Ca^2+^ waves in developing wing discs[46]. This also is accompanied by a reduction in wing size, which correlates recurring calcium waves with growth inhibition[67]. Yet, how perturbed *Gαq* expression impacts wing size and how calcium wave production is specifically involved remain unclear.

Recently, a study reported that inhibition of GPCRs, such as *rickets (rk)*, *methuselah-like 4* (*mthl4*), *CG15744*, and *Stan,* reduce wing size[112]. Here, we confirm that the downregulation of *Gαq* in the wing disc results in smaller wings, independent of the Gal4 driver used. Additionally, we observed a similar size reduction with inhibition of *Gαq* using MS1096-Gal4 and C765-Gal4 drivers. A more severe wing expansion defect was observed in wing discs inhibiting *Gαq* driven by the nubbin-Gal4 driver. Previous work reported that three G-proteins: *Gβ13F*, *Gγ1,* and *Gαs,* are essential for proper wing expansion[113]. Here, our study identified that *Gαq* expression levels also could impact wing expansion. A recent study by Sobala et al. reported that *Gαs*, *Gαq*, and *Gβ13F* are expressed at different stages in the pupal wing disc during pupal wing development[114]. Our study confirms this observation as *Gαq* downregulation leads to defects in pupal wing expansion. Saad et al. reported similar post-expansion defects when GPCRs such as the *methuselah* (*mth*)-like family receptors *mthl6*, *mthl8* and *mthl9* were downregulated in the *Drosophila* wing under nubbin gal4 driver[112]. Future studies are required to identify the GPCRs that function upstream of Gαq to regulate organ growth or expansion phenotypes.

This study highlights the importance of Gαq homeostasis during organ growth. Gαq stimulates IP_3_ production, which activates the IP_3_R channel to release Ca^2+^ from ER[19], [115]–[120]. Our previous experimental studies investigated the emergence of diverse classes of spatiotemporal Ca^2+^ dynamics in both *ex vivo* cultures and *in vivo*[46], [67]. Here, our *in vivo* imaging studies revealed an increase in Ca^2+^ activity and strong intercellular calcium wave (ICW) formation with *Gαq* overexpression (**Fig. 1B**), consistent with past *ex vivo*. This result confirms that Gαq stimulates Ca^2+^ release from ER into the cytoplasm through the activation of its downstream effectors. Our previous studies show that normal physiological levels of calcium signaling promote wing disc growth[67]. Interestingly, further elevating calcium signaling through *Gαq* overexpression reduces wing size, thereby suggesting additional Gαq downstream effectors participating in growth regulation.

To assess whether Gαq*-*mediated Ca^2+^ signaling impacts the size phenotype, we performed genetic interaction experiments by overexpressing *Gαq* and co-expressing RNAi constructs targeting downstream regulators of Ca^2+^ signaling. Surprisingly, we found a statistically significant enhancement in wing size reduction when Ca^2+^ signaling components were inhibited along with *Gαq* overexpression. Thus, we hypothesize that the primary growth-inhibiting activity of Gαq may be through a mechanism independent of Ca^2+^ activity. Moreover, our trichome analysis for *Gαq* overexpression shows that the intervein regions associated with Hh signaling show a different trend in trichome number and density when compared to the overall wing. Our findings from this study imply that Gαq interacts with other growth signaling pathways, thus negatively influencing growth. Together, these results demonstrate the dual action performed by Gαq during wing development, where it stimulates growth through downstream IP_3_R/Ca^2+^ activity and inhibits growth through its interactions with mitogenic pathways independent of Ca^2+^ activity. In addition to *Gαq*, our results show that the depletion of Plc21c levels, which is homologous to mammalian PLCβ, results in decreased adult wing size. This further supports our findings that the depletion of Ca^2+^ signaling components leads to a reduction in wing growth[67]. Future studies are required to investigate the role of Gαq mediated PKC activity in regulating the growth independent of the Ca^2+^ signaling.

Our molecular and transcriptomic analysis investigated the downstream signaling motifs affected in response to *Gαq* perturbations. Our PH3 studies showed an increase in proliferation density for both overexpression and RNAi wing discs. The higher proliferation density observed may be due to wing discs being physiologically younger than the control discs, so they proliferate more rapidly. Our transcriptomic study further confirms this result as we observed an upregulation in genes related to DNA replication for both *Gαq* perturbations. However, the trichome number studies showed a reduction in the overall trichome number for *Gαq*^RNAi^, whereas it did not show any significant change for *Gαq* overexpression. Moreover, the trichome density increased for both perturbations compared to the control and can be attributed to the reduced wing size. As cell size and number are indirect readouts of growth and proliferation, this result establishes that Gαq homeostasis is necessary for maintaining an optimal growth rate and duration of organ growth. Several studies identify the active role of Gαq while promoting growth and proliferation in different systems[14], [15], [121]. For example, Gαq mediates the PI3K activation and the subsequent airway smooth muscle growth[122]. In addition, GPCRs stimulate YAP phosphorylation via Gαq[123], thus regulating Hippo signaling in HEK cells. Taken together, our results demonstrate that Gαq impacts the final wing size by tuning growth and proliferation.

Further, inhibition and overexpression of *Gαq* in the wing disc slightly decrease the total number of cells undergoing apoptosis. Furthermore, our RNA-seq analysis shows both upregulation and downregulation of apoptotic genes for both overexpression and RNAi conditions (**Fig. 6A**). Studies have shown that Gαq regulates cell death both positively and negatively in multiple systems[111], [124], [125]. For example, *Gαq* expression decreases apoptosis in cardiomyocytes through EGFR activation[124]. In addition, Gαq mediated ICWs inhibit excessive apoptosis during epithelial wound healing[126]. Conversely, Gαq activation by GPCR induces apoptosis by reducing PKC-dependent AKT activity in prostate cancer cells[125]. Collectively, these studies provide evidence that Gαq either stimulates or inhibits apoptosis. Consistent with these studies, our transcriptomic study shows that *Gαq* perturbations resulted in dysregulation of the apoptotic genes. However, our results show that inhibition of apoptosis along with *Gαq* overexpression could not rescue the wing size phenotype, thereby suggesting a minimal role of apoptosis in regulating Gαq*-*mediated growth reduction.

Our transcriptomic study revealed that *Gαq* overexpression activates stress response genes, which include heat shock proteins and serine proteases. The expression of serine proteases is increased as serpins, which inhibit serine proteases, were downregulated in our RNA-seq data. In addition, our findings show that JNK components, including *dilp8*, *upd3,* and *chinmo* were significantly upregulated. JNK signaling induces stress and immune response[127]–[130] and initiates compensatory proliferation[130]. Importantly, JNK signaling induces *dilp8* expression in the wing disc in response to tissue damage or stress[83]. Further, increased Dilp8 levels communicate with the brain to inhibit the ecdysone synthesis, thereby slowing growth[83], [131]. Consistent with these reports, we found that *Gαq* overexpression decreased the expression of 20E-dependent genes. Moreover, our developmental assay confirms this result as we observed a delay in larval to pupal transition for flies expressing *Gαq* in the wing disc. This developmental delay is strikingly similar to that observed when *dilp-8* is overexpressed in the wing disc[83], [84]. Studies have shown that different *Gα* subtypes such as *Gαo, Gα_16_* induce mitogenic activity through activation of JAK/STAT signaling[38], [132], [133]. For instance, the constitutively active form of *Gαo* induces activation of STAT3 in NIH-3T3 cells and subsequent cell transformation[133, p. 3]. In addition, Heasley et al. have shown that expression of the constitutively active form of *Gα_16_* inhibits growth and activates JNK signaling in small lung carcinoma cell lines (SLC)[39]. By contrast, sustained activation of Gαq signaling induces cell death in GABAergic medium spiny neurons via activation of JNK signaling[134]. These studies indicate the active role of JAK/STAT signaling in Gαq mediated growth and apoptosis. Cumulatively, our results demonstrate that *Gαq* overexpression regulates JNK signaling genes, which might possibly regulate apoptosis and proliferation, thereby affecting the final wing size. While many studies have investigated the mechanisms underlying the transmission of proliferative signals through GPCR stimulation *in vitro*, further research is required to elucidate how G proteins encode mitogenic signals *in vivo*[14], [135].

Next, our RNA seq analysis of *Gαq*^RNAi^ indicates that the nuclear hormone receptor-related genes were affected (**Fig. 6E** **& 7B**). Our analysis indicated that DNA replication was enriched among the upregulated genes. Moreover, the downregulated genes were primarily involved in biological processes related to cuticle development, molting cycle, taxis, and transmembrane transport. Interestingly, we observed downregulation in genes mediating ecdysone response (**Fig. 6E**), thereby resulting in the downregulation of ecdysone-induced genes (**Additional File 1, Fig. S13 and Fig. S15**). Specifically, the *Ecdysone-induced genes 71E* family (*Eig71E*), which are regulated by Ecdysone-induced primary response genes, were downregulated (**Additional File 1, Fig. S13, Additional File 3)**. A further finding was an increase in Dilp-8 levels, which may explain the inhibition of genes related to ecdysone signaling. Our developmental assay further confirmed this as we observed an approximate twelve-hour delay in the pupariation of flies expressing *Gαq*^RNAi^. Interestingly, we identified a similar trend in gene expression profile for both *Gαq*^RNAi^ and *Gαq* overexpression conditions. However, the extent of gene activation or suppression is statistically less in *Gαq*^RNAi^ when compared to the *Gαq* overexpression. These results suggest a common compensatory mechanism in a cell operating in response to dysregulation of Gαq homeostasis. Moreover, our trichome analysis of *Gαq*^RNAi^ showed a reduction in the cell number, thereby indicating that the reduction in the wing disc growth is caused by the lack of calcium signaling. Collectively, our RNA seq results demonstrated that disruption to Gαq homeostasis reduced growth and increased developmental time. Further, the increased developmental time caused by *Gαq* overexpression occurs through Ca^2+^ activity. Significantly, we observed that inhibition of Ca^2+^ signaling through Ip_3_R knockdown rescues the development delay caused by *Gαq* overexpression. However, the knockdown of Ip_3_R along with *Gαq* overexpression reduced wing size, thereby implying that Ca^2+^ signaling promotes proliferation in the wing disc by regulating the developmental time.

Importantly, our RNA seq data revealed a reduction in the expression of serine protease inhibitors, thus resulting in an increase in the expression of serine proteases, including Jonah genes, which are key proteolytic enzymes (**Additional File 2)**. Moreover, serine protease cascades are known to elicit immune responses by inducing proteolytic cleavage of *spz*[92]. Our enrichment analysis investigating the core signaling pathways indicated that the nuclear hormone receptor, Toll signaling, and JAK/STAT were significantly affected in the *Gαq* overexpression condition. This result confirms that disruption to Gαq homeostasis parallels signals related to tissue damage response, thereby eliciting stress and immune response. Several studies have reported the role of G proteins in mediating immune response and autoimmunity[136]–[138]. For example, Zhu. et al. reported that XLG2, a functional *Gα* protein, positively regulates immune response to provide disease resistance in plants[139]. Interestingly, studies have shown that the activation of immune signaling occurs at the expense of growth[89]. Further investigations are needed to explore the crosstalk mechanisms between Gαq and immune regulation during organ growth.

Of note, our transcriptomic analysis of *Gαq* OE shows a strong upregulation in the orphan GPCR Methuselah-like 8 (mthl8) with a striking log2 fold change of 11.5 (**Additional File 4**). *Mthl-8* has been shown to impact wing expansion[140]. In addition, the family of Methuselah genes is known to be involved in longevity and stress resistance[141]. Recent work in Lepidoptera suggests that GPCRs with homology to the Methuselah family of receptors may play important, unresolved roles in mediating the key hormone ecdysone[63], [142]. Importantly, it has been shown that *mthl-8* is a potential target of the JNK signaling in the *Drosophila* eye disc[143]. Collectively, our results imply a crosstalk between Gαq and GPCRs such as mthl-8. Further investigations are needed to elucidate the role of G proteins in regulating the expression of signal transducers such as GPCRs.

Moreover, studies are needed to clarify the subcellular location and functional activity of Gαq and related GPCRs. Recently, GPCRs have been found to be active not only in the outer cell membrane but associated with the inner membranes of organelles within the cell[144], [145]. Also, further investigation is needed to map out key molecular players activating Gαq-mediated ICWs *in vivo*. For instance, recent studies have indicated that growth-blocking peptide (Gbps) binds to Mthl10 receptors and triggers Gαq mediated ICWs in the *Drosophila* wing disc[48]. Understanding the growth regulatory mechanisms acting through G-proteins may lead to broadly applicable insights into cancer therapeutics and drug development[141], [146].

## Conclusions

In summary, our study has decoded the downstream impacts of perturbing *Gαq* expression. Gαq activity is sufficient to stimulate intercellular calcium waves and is a key regulator of Ca^2+^ activity during development and wound healing. Our study characterized the multiple growth modules impacted by *Gαq* perturbations and their roles in facilitating organ growth control during development. An intriguing finding is that there exist similar changes in gene expression for *Gαq*^RNAi^ and *Gαq* OE conditions, which suggests that homeostasis of *Gαq* expression is necessary for proper growth and development. Overall, our findings from this work suggest that a central functional role of Gαq*-*mediated Ca^2+^ signaling is to coordinate the growth status of peripheral organs to the brain by upregulating Dilp8, thereby mediating the “crosstalk” between developing organs and “the control center” of the brain (**Fig. 6E**). Thus, tissue wide Ca^2+^ waves generated by Gαq activity coordinates the response to wounding or other stressors to extend the developmental time and reduce growth.

## Materials and methods

### Fly stocks

*Drosophila* stocks were grown on standard laboratory cornmeal food. All crosses were set up at 25°C. The following stocks were used for the experiment: C765-Gal4 y[1] w[*]; P{w[+mW.hs]=GawB}C-765 (BDSC:36523), nubbin-GAL4, UAS-Dcr-2 (BDSC:25754), MS1096- GAL4, UAS-Dcr-2 (BDSC:25706), UAS-Gαq (BDSC:30734), UAS-Galpha49B RNAi (BDSC:36775), w; Sco/CyO; Dr/TM6B, Tb (gift of Carthew Lab), w; Sco/CyO; MKRS/TM6B, Tb (gift from Carthew lab), w; UAS-Gαq/CyO; C765-Gal4/TM6B,Tb (generated in this study), w; UAS- Gαq,C765-Gal4/Cyo-TM6B,Tb (generated in this study), UAS-Itpr RNAi (BDSC:25937), UAS-p35 (BDSC #6298), UAS-Plc21C RNAi (BDSC: 33719), UAS-sl RNAi (BDSC:32906), UAS-Rya-r44F RNAi (BDSC:31540), nub-GAL4, UAS-GCaMP6f/CyO (recombinant line made in Zartman Lab).

### Immunohistochemistry and imaging of fixed samples

The imaginal wing discs were dissected in Phosphate Buffered Saline (PBS) (#P5368, Millipore-Sigma). Following the dissection of the wing discs, they were fixed in 4% PFA for 20 minutes at room temperature, followed by three 10-minute washes in PBT (0.03% Triton X-100 (#T9284, Sigma-Aldrich) in PBS). Next, discs were blocked in 5% Normal Goat Serum (NGS) (#NC9660079; Jackson Immuno Research Laboratories, West Grove, PA) for 30 minutes and followed by overnight incubation with primary antibodies at 4°C. The following primary antibodies were used: PH3 (#9701S, Cell Signaling), diluted to 1:800 in 5% NGS, and DCP-1 (#9578S; Cell Signaling), diluted to 1:100 in 5% NGS. Incubation of the primary antibodies was followed by three 15-minute washes with PBT and subsequent incubation with Alexa Fluor-conjugated secondary antibodies (1:500 in 5%NGS) and DAPI (# D9542; Millipore Sigma) (1:1000) for 2-3 hours at room temperature in the dark. After this, wing discs were washed with PBT overnight at 4°C, followed by two 15-minute washes at room temperature. Discs were next mounted in Vectashield (# H- 1000, Vector Laboratories) on a 24×66 mm coverslip and covered by a 22×22 coverslip, and sealed with clear nail polish by the sides. Images of mounted wing discs were obtained using a Nikon Eclipse Ti confocal microscope with a Yokogawa spinning disk (Tokyo, Japan) at a magnification of 40x (oil objective, NA 1.30).

### In vivo imaging setup

Third instar wandering larvae were collected and rinsed in deionized water prior to imaging. A coverslip was used to image the larvae after they had been dried and adhered to a cover with Scotch tape. The larvae were attached with their spiracles facing the coverslip to align the wing discs toward the microscope. An EVOS FL Auto microscope was used to image the larvae at a magnification of 20x for 20 minutes. The images were taken every 15 seconds.

### RNA extraction

RNA was isolated using RNeasy Mini Kit (#74104; QIAGEN), using a standard procedure with DNase on column digestion (RNase-free DNase set. #79254, QIAGEN). RNA was eluted in RNase DNase-free water provided by the RNeasy Mini Kit. Samples from multiple days of dissection were pooled such that each biological replica would have 79 WDs. To assess the RNA quantity and quality, we performed QC using Qubit and Agilent Bioanalyzer and polled samples from multiple days of RNA isolation to achieve at least 50ng of RNA per sample or higher.

### Sequencing

RNASeq libraries were prepared and sequenced across one lane of an Illumina NextSeq v2.5 (Mid Output 150 cycle) flow cell. Each library was prepared using the NEBNext Ultra II Directional RNA. Library Prep kit and the NEB mRNA Magnetic Isolation Module. We performed QC and quantitation on the library pool using the Qubit dsDNA Agilent Bioanalyzer DNA. High Sensitivity Chip, and Kapa Illumina Library Quantification qPCR assays. The sequencing form was paired-end 75bp. Base calling was done by Illumina Real Time Analysis (RTA) v2 software.

### Processing of sequencing data

We trimmed the raw sequences of adapters with Trimmomatic version 0.39[147] and assessed for quality with FastQC v 0.11.8[148]. Trimmed sequences were aligned to the *Drosophila* genome, and Ensembl built *Drosophila melanogaster.*BDGP6.32.104.gtf, Berkeley *Drosophila* Genome Project (BDGP, Release 6, Aug. 2014), using Dmel_Release_6.01 version annotations and HISAT2 version 2.1.0[149]. SAMtools version 1.9 was used to sort the corresponding alignments[150]. Read counts were generated with HTSeq-count version 0.11.2[151]. Subsequent statistics were completed in R (R Core Team, 2014), implementing the edgeR library version 3.36.0[152]–[155]. Using the Ensembl version of BioMart, Gene names and GO terms were identified[156].

### Quantification of adult wings and statistics

Using ImageJ, we measured the total area of the wings. The wing margin was traced by following veins L1 and L5, and the hinge region was excluded from the size analysis. All statistical analyses were performed using R and Excel. Student t-tests were performed to assess the statistical significance of our comparisons. P-value, standard deviation, and sample size (n) are provided in each figure and legend.

### Developmental timing assay

Virgins were 1-3 days old when mated with males. Crosses were mated 24 h prior and transferred to a vial for egg collection. Fertilized eggs from each cross were collected for eight hours in a fresh vial with corn meal food and then transferred to a new vial. After egg laying, pupae were manually scored in 12-hour intervals for ten days. For C765>Gαq + >RyR^RNAI^ and C765>Gαq + >Ip_3_R^RNAi^, pupae were manually counted in 24-hour intervals for approximately 6 to 7 days.

## Abbreviations

AkhR: Adipokinetic Receptor
chinmo: chronologically incorrect morphogenesis
DAG: diacylglycerol
DAPI: 4′,6-Diamidino-2-phenylindole dihydrochloride
DCP: Cleaved Drosophila Dcp-1 (Asp215) Antibody
DCP-1: death caspase-1
Dpp: Decapentaplegic
DUOX: Dual oxidase
Eig: Ecdysone-induced gene
ER: endoplasmic reticulum
ERK: extra-cellular-signal-regulated protein kinase
GDP: guanosine diphosphate
GO: Gene Ontology
GPCRs: G protein-coupled receptors
GTP: guanosine triphosphate
Gαq O.E.: G alpha q Overexpression
Gαq RNAi: G alpha q RNAi
Gαq: G alpha q
Hh: Hedgehog
Hr: Hormone receptor
Hsp: Heat shock protein
ICWs: intracellular calcium waves
Im33: Immune-induced molecule 33
IP3: inositol trisphosphate
IP3R: IP3 Receptor
JNK: Jun-N-terminal-Kinase
Lys: Lysozyme
MAPK: mitogen-activated protein kinase
mthl: methuselah-like
NGS: Normal Goat Serum
Oatp74D: Organic anion transporting polypeptide 74D
PBS: Phosphate-buffered saline
Pdk1: Phosphoinositide-dependent kinase 1
PH3: Phospho-Histone H3 (Ser10) Antibody
PKC: protein kinase C
PLC-β: phospholipase Cβ
rk: rickets
RNA: Ribonucleic acid
RNAi: RNA interference
Serpin, Spn: Serin Protease Inhibitor
Spz: Spatzle
upd3: unpaired3
Wg: wingless

## Acknowledgments

The work in this paper was supported by NIH Grant R35GM124935, NSF Award CBET-1553826, and EMBRIO Institute, contract #2120200, a National Science Foundation (NSF) Biology Integration Institute. Notre Dame Genomics & Bioinformatics Core Facility provided services related to RNA sequencing. The authors would also like to thank members of the Zartman lab for helpful discussions. We would like to thank Giorgia Giordano for her help in the project.

## Author contributions

VVK performed experiments, analyzed data, and wrote the paper. DKS conceived the study, performed experiments, analyzed data, and wrote the paper. MU performed experiments, analyzed data, and wrote the paper. DG and NK performed experiments and analyzed data. JL performed the RNA seq analysis. JJZ conceived the study, analyzed data, wrote the paper, and supervised the study.

## Data Availability

Differential gene expression analysis data obtained from our RNA sequencing study is available in Additional File 3 and Additional File 4. The RNA sequencing raw data and the wing image data obtained during the current study is available from the corresponding author on reasonable request.

## Conflict of interests

The authors declare no conflicts of interest.

## List of Additional Files

- Additional File 1
  - Document (.pdf)
  - Supplementary text
  - This file includes the figure for the trichome analysis of the intervein areas. It also includes the figures of the results obtained from the GO enrichment analysis of the differentially expressed genes. In addition, it contains supplementary text for the methods used to analyze the wings and the methods used to plot the GO enrichment results.
- Additional File 2
  - spreadsheet (.csv)
  - Fold changes of differentially expressed genes associated with immune-related GO terms.
  - This file contains the fold changes of differentially expressed genes involved in immune response in response to both *Gαq* perturbations.
- Additional File 3
  - spreadsheet (.csv)
  - Differentially expressed genes between C765>Ryr^RNAi^ and *C765>Gαq^RNAi^*
  - This file contains the results of the differential gene expression analyses of the RNA sequencing data for *C765>Ryr^RNAi^* vs. *C765>Gαq^RNAi^*.
- Additional File 4
  - spreadsheet (.csv)
  - Differentially expressed genes between C765>Ryr^RNAi^ and *C765>Gαq*
  - This file contains the results of the differential gene expression analyses of the RNA sequencing data for *C765>Ryr^RNAi^* vs. *C765>Gαq*

## Notes

### Competing Interest Statement

The authors have declared no competing interest.

