## Additional File 1 for "The multimodal action of G alpha q in coordinating growth and homeostasis in the *Drosophila* wing imaginal disc"

<u>**Supplementary text**</u>

#### Contents


1) **SI Methods:** Methods for RNA seq analysis and wing area measurements.
2) **Figure S1**: Disruption of Gαq homeostasis affects the cell number and cell density in the intervein regions.

**Gene expression analysis of wing discs expressing *Gαq***

3) **Figure S2:** Gene ontology enrichment analysis of differentially expressed genes in wing discs expressing *Gαq* (*C765>GαqOE*).
4) **Figure S3**: Gene ontology enrichment analysis of the genes upregulated in wing discs expressing *Gαq* under *C765-Gal4* driver (FC ≥ 1.5).
5) **Figure S4:** Network representation of the top GO terms enriched among the significantly upregulated genes in wing discs expressing *Gαq* with the *C765-Gal4* driver (FC ≥ 1.5).
6) **Figure S5:** Overexpression of *Gαq* in the wing disc with the *C765* driver results in the upregulation of genes associated with serine proteases and cuticle proteins.
7) **Figure S6:** Gene ontology enrichment analysis of the genes downregulated in wing discs expressing *Gαq* with *C765-Gal4* driver (FC ≤ 0.65).
8) **Figure S7:** Network representation of the top GO terms enriched among the significantly downregulated genes in wing discs expressing *Gαq* under *C765-Gal4* driver (FC ≤ 0.65).
9) **Figure S8**: Overexpression of *Gαq* in the wing disc with the *C765-Gal4* driver downregulated genes involved in the cellular response to ecdysone, extracellular matrix, and negative regulation of the metabolic process.

**Gene expression analysis of wing discs expressing $G\alpha q^{RNAi}$**

10) **Figure S9**: Gene ontology enrichment analysis of differentially expressed genes in wing discs expressing $G\alpha q^{RNAi}$ ($C765>G\alpha q^{RNAi}$).
11) **Figure S10**: Gene Ontology enrichment analysis of upregulated genes for $G\alpha q^{RNAi}$ expression in wing discs ($C765>G\alpha q^{RNAi}$) with FC ≥ 1.5.

12) **<u>Figure S11</u>**: Expression of *Gαq*$^{RNAi}$ in the wing disc under *C765* driver upregulated genes associated with peptidase activity and DNA replication.

13) **<u>Figure S12</u>**: Gene Ontology enrichment analysis of downregulated genes for *Gαq*$^{RNAi}$ expression in wing discs (*C765>Gαq*$^{RNAi}$) with FC ≤ 0.65.

14) **<u>Figure S13</u>**: Expression of *Gαq*$^{RNAi}$ in the wing disc under *C765* driver downregulated genes involved in programmed cell death, defense response, and cell projection assembly.

**Polar plots showing the fold changes of major signaling components for *Gαq* perturbations.**

15) **<u>Figure S14:</u>** Circular plot illustrating the fold change values of core signaling pathway genes that were differentially expressed in wing discs over expressing Gαq with the C765-Gal4 driver.

16) **<u>Figure S15:</u>** Circular plot illustrating the fold change values of core signaling pathway genes that were differentially expressed in wing discs expressing GαqRNAi with the C765-Gal4 driver.

17) **<u>SI References</u>**

### SI Methods

### Methods for RNA seq analysis and wing area measurements.

#### Analysis of RNA seq data

The read count matrix was processed using edgeR to obtain the differential gene expression data. For differential gene expression analysis, we followed an approach similar to that of Ghosh et al.[1]. Using the clusterprofiler[2], we conducted Gene Ontology enrichment analysis among the differentially expressed genes. To plot the measurements obtained from the Gene ontology enrichment analysis, we used the custom dotplot, emaplot, and cnet plot functions of the cluster profiler package. To perform pathway enrichment analysis, we first obtained the components of the major signaling pathways from GLAD[3]. We then performed a hypergeometric test on the differential gene expression data to determine the enrichment of the core signaling pathways. A heatmap2 package was used to plot enrichment scores and adjusted p-values obtained from the analysis. Components for the GO terms: cell death, cuticle proteins, and ecdysone response were obtained from the fly base. Fold changes for the components of each GO term were imported to R. A heat map was plotted using the heatmap2 package to plot the genes having significant fold change. Genes that were absent in the differential gene expression data or did not show a significant fold change were plotted as zero in the heat map plot. For bubble plots, we imported the differential gene expression data into Tableau and plotted the log fold change values using the bubble plot built-in graph.

#### Analysis of trichome number and trichome density

We used Ilastik[4] to generate segmentation masks for the wing images. Next, wing images and the segmentation masks were imported to MAPPER[5] to quantify the wings. The trichome number and area for each intervein region were obtained and further processed in R to calculate the trichome density.

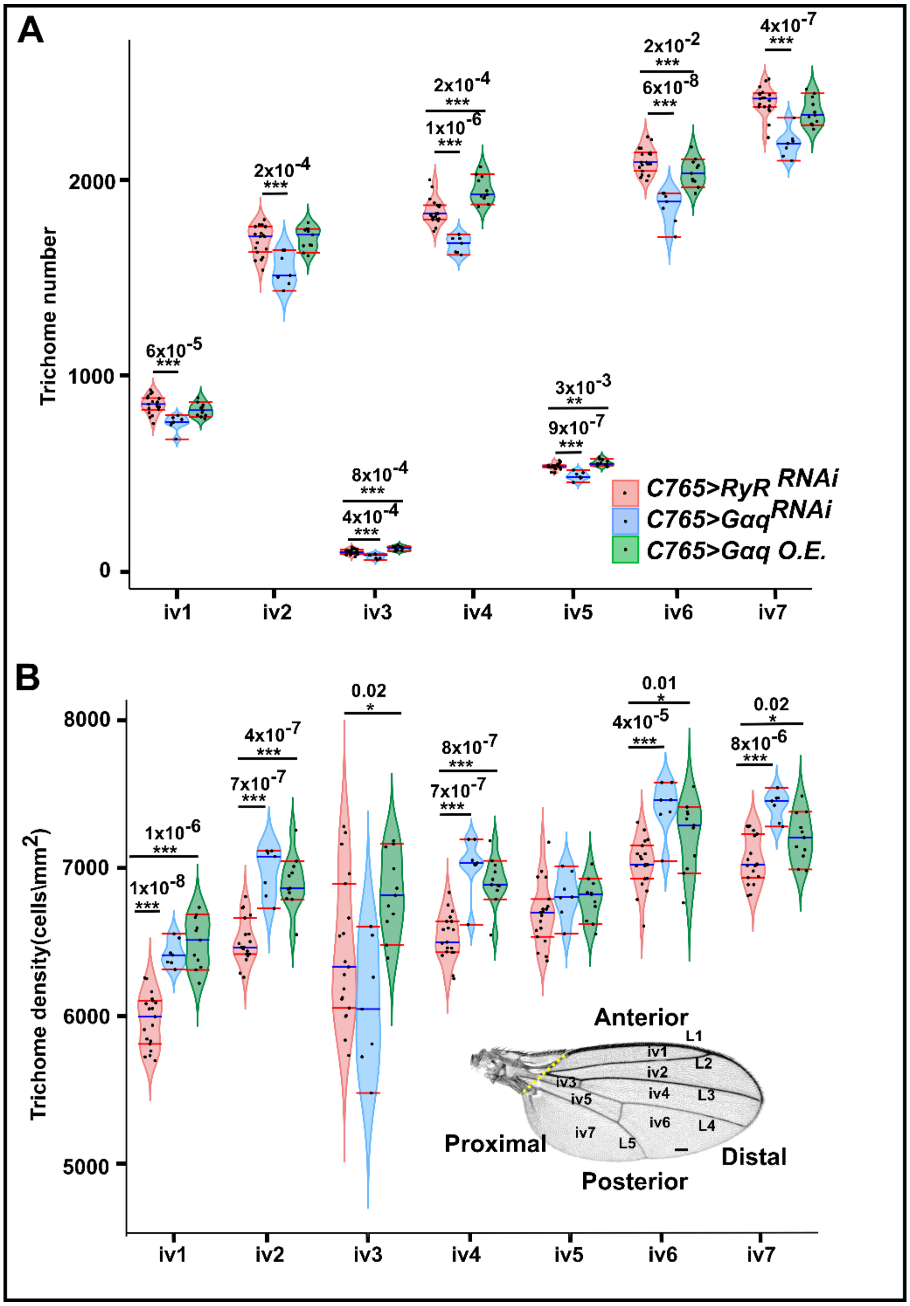


**Figure S1**: **Disruption of Gαq homeostasis affects the cell number and cell density in the intervein regions. A)** Violin plot comparing the trichrome number at each intervein region of the wings expressing *Ryr*$^{RNAi}$ (Control), *Gαq*$^{RNAi}$, and *Gαq* under the *C765-Gal4* driver. The trichome number is reduced for each intervein region in the wings expressing *Gαq*$^{RNAi}$. *Gαq* overexpression increased the trichome number in intervein regions 2, 3, 4, and 5. B) Plot showing the trichome density in each intervein region of the wings expressing $Ryr^{RNAi}$ (Control), $G\alpha q^{RNAi}$, and *GαqOE*. *GαqOE* and $G\alpha q^{RNAi}$ showed increased trichome density for the intervein regions 1, 2, 4, 6, and 7.

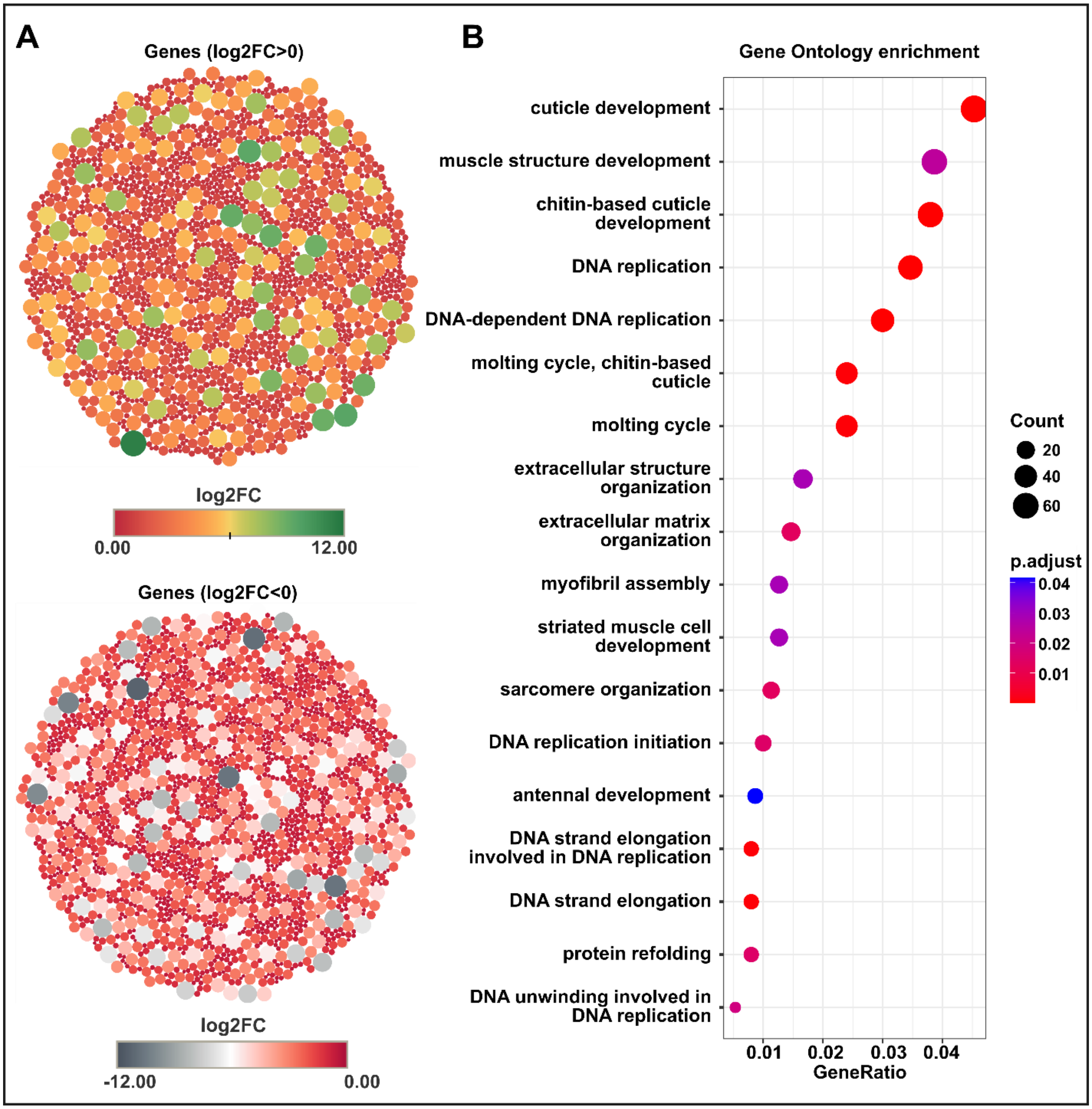


**Figure S2: Gene ontology enrichment analysis of differentially expressed genes in wing discs expressing *Gαq* (*C765>Gαq OE*). A)** Bubble plot showing an overview of the genes that have a significant fold change value in wing discs expressing *Gαq* under *C765-Gal4* driver. The top plot depicts the upregulated genes with a statistically significant fold change value, while the bottom plot depicts the downregulated genes with a statistically significant fold change value. Each bubble represents a gene. The bubble size represents the magnitude of the absolute fold change value, whereas color represents the fold change value. **B)** Dot plot showing the top significant enriched Gene Ontology terms (adjusted p-value ≤ 0.05) for the genes with a significant fold change (-0.6 ≥ log2FC ≥ 0.6). Gene ratio is a ratio of genes with statistically significant differential expression in this dataset versus a total number of genes representing each of the top GO terms.

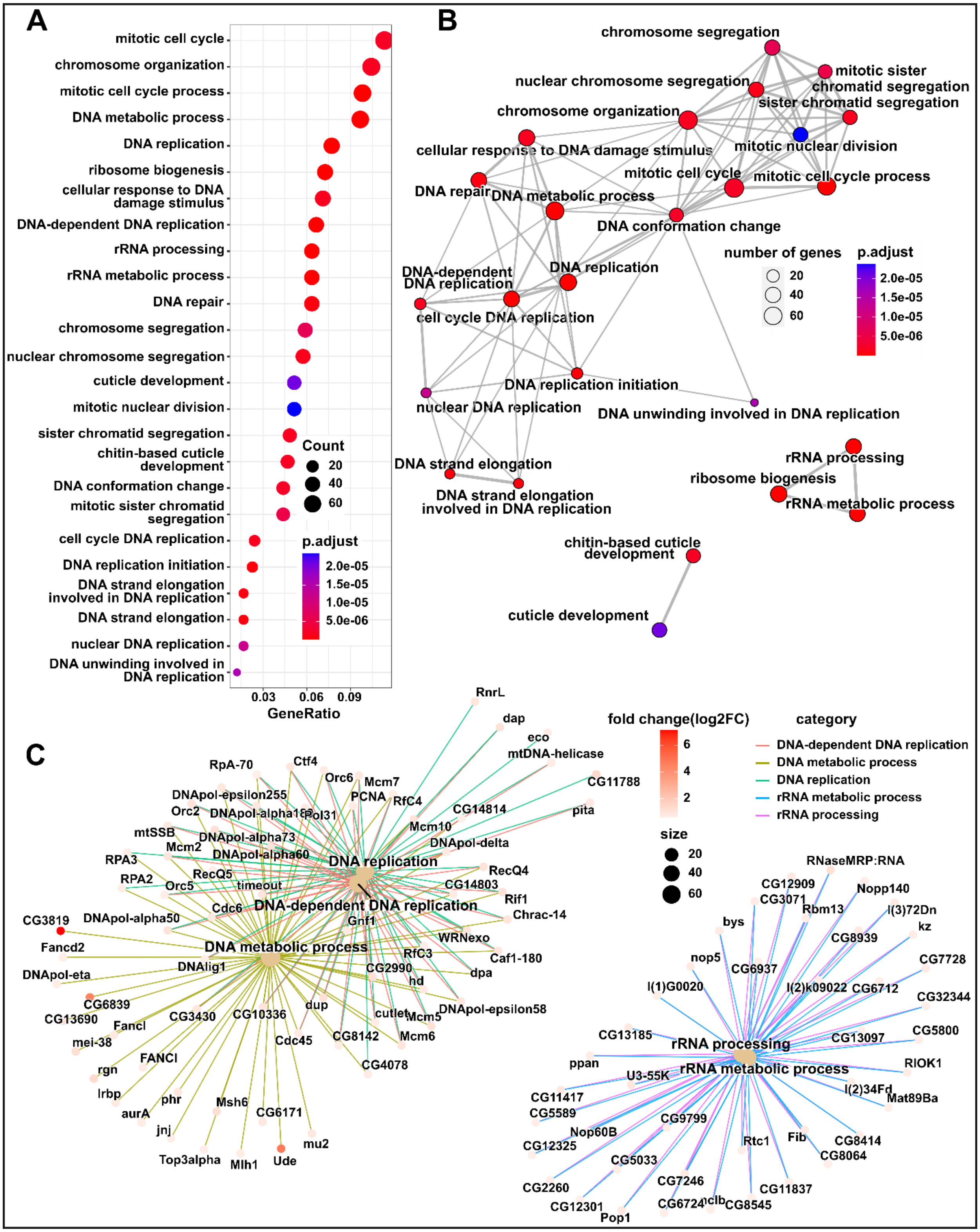
A
mitotic cell cycle
chromosome organization
mitotic cell cycle process
DNA metabolic process
DNA replication
ribosome biogenesis
cellular response to DNA damage stimulus
DNA-dependent DNA replication
rRNA processing
rRNA metabolic process
DNA repair
chromosome segregation
nuclear chromosome segregation
cuticle development
mitotic nuclear division
sister chromatid segregation
chitin-based cuticle development
DNA conformation change
mitotic sister chromatid segregation
cell cycle DNA replication
DNA replication initiation
DNA strand elongation involved in DNA replication
DNA strand elongation
nuclear DNA replication
DNA unwinding involved in DNA replication
Count
20
40
60
p.adjust
2.0e-05
1.5e-05
1.0e-05
5.0e-06
0.03
0.06
0.09
GeneRatio
B
chromosome segregation
mitotic sister chromatid segregation
nuclear chromosome segregation
sister chromatid segregation
chromosome organization
cellular response to DNA damage stimulus
mitotic nuclear division
mitotic cell cycle
mitotic cell cycle process
DNA repair
DNA metabolic process
DNA conformation change
number of genes
20
40
60
p.adjust
2.0e-05
1.5e-05
1.0e-05
5.0e-06
DNA-dependent DNA replication
DNA replication
cell cycle DNA replication
DNA replication initiation
nuclear DNA replication
DNA unwinding involved in DNA replication
DNA strand elongation
DNA strand elongation involved in DNA replication
rRNA processing
ribosome biogenesis
rRNA metabolic process
chitin-based cuticle development
cuticle development
C
fold change(log2FC)
6
4
2
category
DNA-dependent DNA replication
DNA metabolic process
DNA replication
rRNA metabolic process
rRNA processing
size
20
40
60
DNA replication
DNA-dependent DNA replication
DNA metabolic process
rRNA processing
rRNA metabolic process
RnrL
dap
eco
mtDNA-helicase
CG11788
pita
RpA-70
Ctf4
Orc6
Mcm7
PCNA
RfC4
DNApol-epsilon255
Orc2
DNApol-alpha180
Pol31
CG14814
mtSSB
Mcm10
DNApol-alpha73
Mcm2
DNApol-delta
DNApol-alpha60
RPA3
RecQ4
RecQ5
RPA2
Orc5
timeout
CG14803
Rif1
Cdc6
Chrac-14
Gnf1
CG3819
DNApol-alpha50
WRNexo
Fancd2
Caf1-180
RfC3
DNApol-eta
DNAlig1
CG2990
dpa
hd
CG6839
DNApol-epsilon58
CG13690
CG3430
CG10336
dup
cutlet
Mcm5
FancI
mei-38
Cdc45
CG8142
Mcm6
FANCI
rgn
CG4078
Irbp
phr
Msh6
CG6171
aurA
jnj
mu2
Top3alpha
Mlh1
Ude
RNaseMRP:RNA
CG12909
Nopp140
CG3071
Rbm13
l(3)72Dn
bys
kz
CG8939
nop5
CG7728
CG6937
l(1)G0020
l(2)k09022
CG6712
CG32344
CG5800
CG13185
CG13097
ppan
RIOK1
U3-55K
l(2)34Fd
Mat89Ba
CG11417
CG5589
CG9799
Nop60B
CG8414
CG12325
Rtc1
Fib
CG8064
CG5033
CG7246
CG2260
CG11837
CG12301
CG6724
nclb
CG8545
Pop1

**Figure S3: Gene ontology enrichment analysis of the genes upregulated in wing discs expressing *Gαq* under *C765-Gal4* driver (log FC ≥ 0.6). A)** A dot plot displaying the top 25 GO terms that were significantly (adjusted p-value ≤ 0.05) enriched among the statistically significantly upregulated genes (log2FC ≥ 0.6). The color of the dot represents the adjusted p-value, and the size represents the number of genes upregulated in the respective category. **B)** Emaplot illustrating the network representation of the top 25 significantly enriched GO terms. GO terms that have mutual overlap are clustered together in terms of similarity. The edges between GO terms are displayed if they share a certain degree of similarity (the default threshold is 0.2). The edges become shorter and thicker as the degree of similarity increases. Each GO term is represented by a node. The color of the node represents the adjusted p-value, and size represents the number of genes that were statistically significantly upregulated in the GO term. C) A cnet plot showing the top five enriched GO terms mapped to the corresponding genes which were statistically significantly upregulated. Each leaf node corresponds to a gene, and the node's color depicts the log fold change.

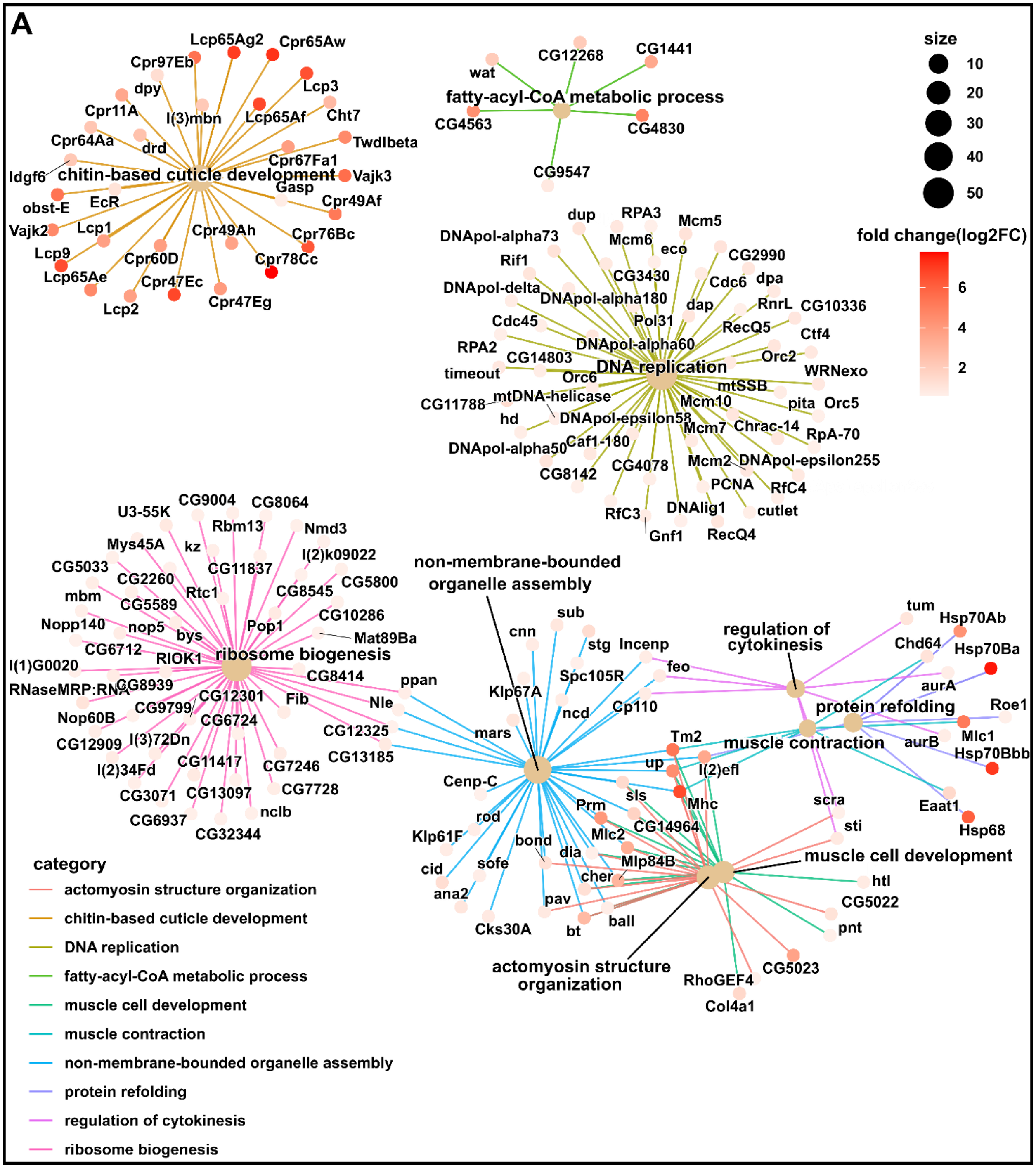


**Figure S4: Network representation of the top GO terms enriched among the significantly upregulated genes in wing discs expressing *Gαq* under *C765-Gal4* driver (log2FC ≥ 0.6). A)** A cnet plot showing the top 10 enriched GO terms after removing the redundant enriched GO terms and the corresponding significantly upregulated genes. Each leaf node corresponds to a gene, and the node's color depicts the log fold change.

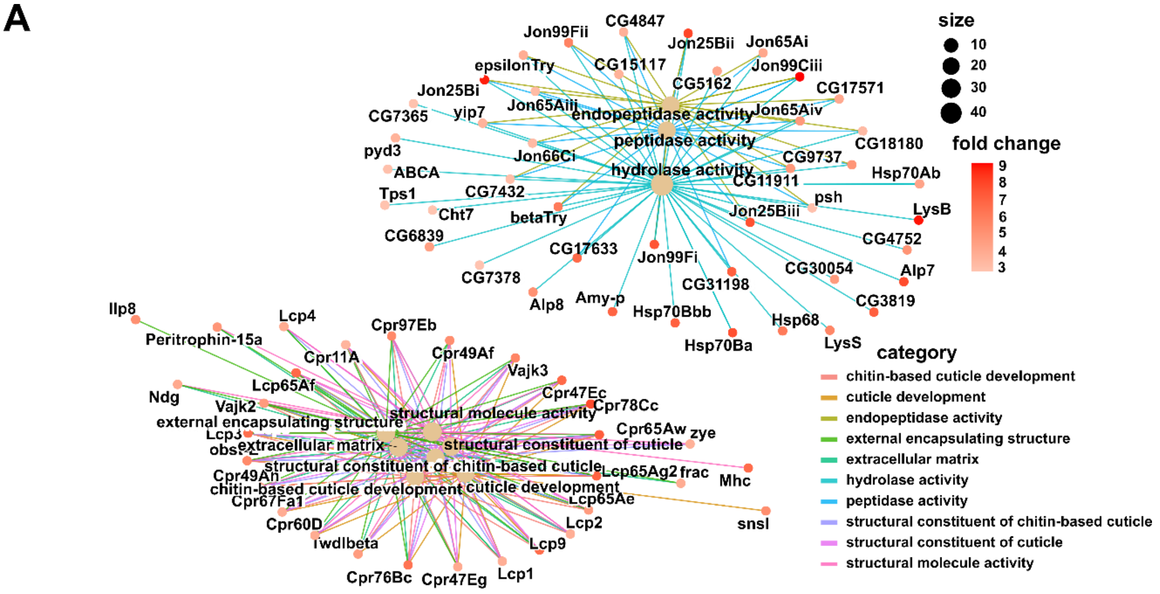


**Figure S5: Overexpression of *Gαq* in the wing disc under the *C765* driver results in the upregulation of genes associated with serine proteases and cuticle proteins**. A) A network representation of the GO terms related to peptidase activity, cuticle synthesis, and the corresponding genes that were upregulated. Each leaf node corresponds to a gene, and the node's color indicates the log fold change.

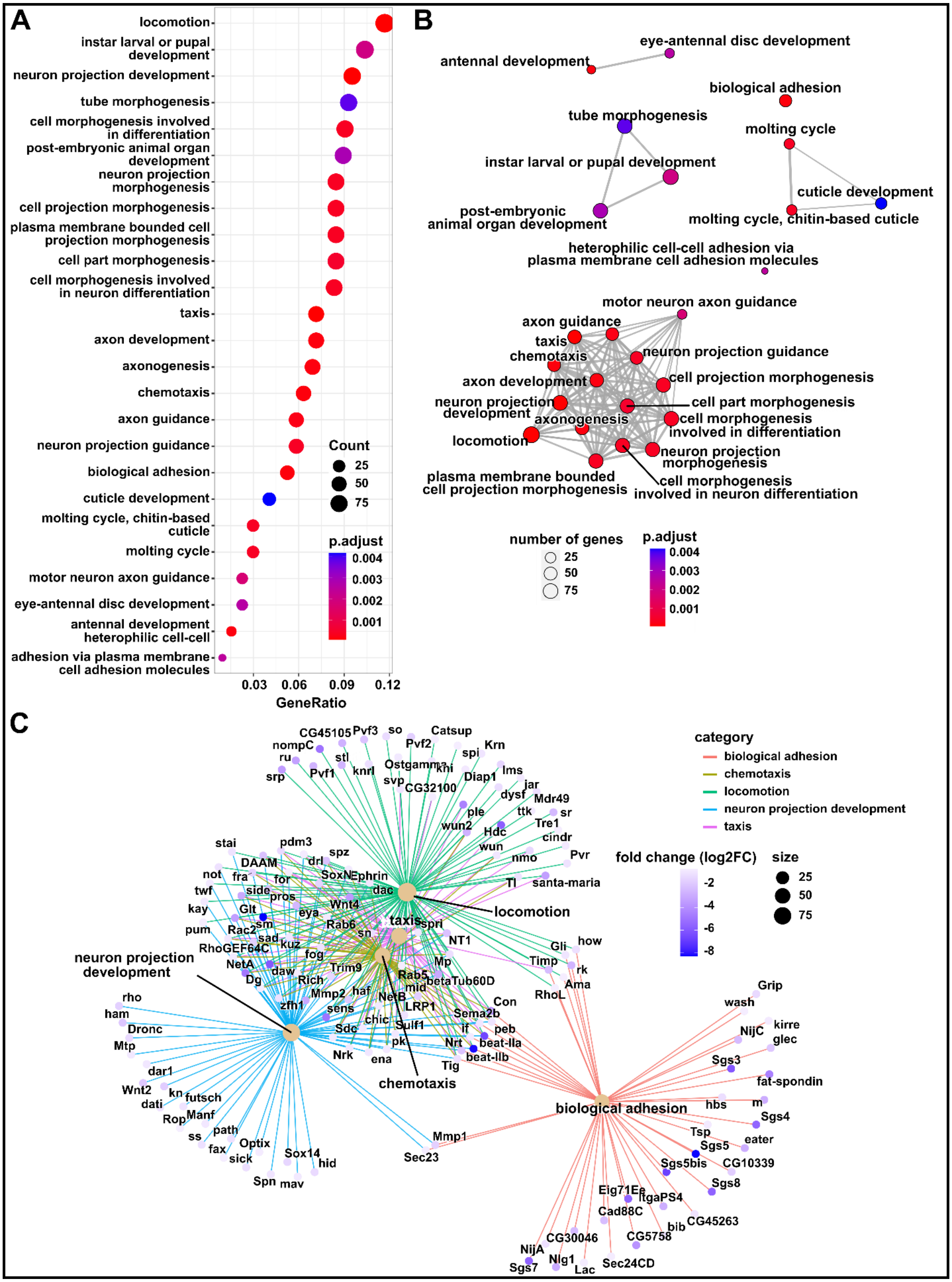
A
locomotion
instar larval or pupal development
neuron projection development
tube morphogenesis
cell morphogenesis involved in differentiation
post-embryonic animal organ development
neuron projection morphogenesis
cell projection morphogenesis
plasma membrane bounded cell projection morphogenesis
cell part morphogenesis
cell morphogenesis involved in neuron differentiation
taxis
axon development
axonogenesis
chemotaxis
axon guidance
neuron projection guidance
biological adhesion
cuticle development
molting cycle, chitin-based cuticle
molting cycle
motor neuron axon guidance
eye-antennal disc development
antennal development
heterophilic cell-cell adhesion via plasma membrane cell adhesion molecules
Count
25
50
75
p.adjust
0.004
0.003
0.002
0.001
0.03
0.06
0.09
0.12
GeneRatio
B
eye-antennal disc development
antennal development
biological adhesion
tube morphogenesis
molting cycle
instar larval or pupal development
cuticle development
post-embryonic animal organ development
molting cycle, chitin-based cuticle
heterophilic cell-cell adhesion via plasma membrane cell adhesion molecules
motor neuron axon guidance
axon guidance
taxis
neuron projection guidance
chemotaxis
axon development
cell projection morphogenesis
neuron projection development
cell part morphogenesis
axonogenesis
cell morphogenesis involved in differentiation
locomotion
neuron projection morphogenesis
plasma membrane bounded cell projection morphogenesis
cell morphogenesis involved in neuron differentiation
number of genes
25
50
75
p.adjust
0.004
0.003
0.002
0.001
C
category
biological adhesion
chemotaxis
locomotion
neuron projection development
taxis
fold change (log2FC)
-2
-4
-6
-8
size
25
50
75
locomotion
taxis
neuron projection development
chemotaxis
biological adhesion
CG45105 Pvf3 so Catsup
nompC Pvf2 Krn
ru stl spi
srp Pvf1 knrl Ostgamma khi Diap1 lms
svp CG32100 jar
dysf Mdr49
ple ttk sr
wun2 Hdc Tre1
cindr
wun
nmo Pvr
stai pdm3 spz
DAAM drl
not fra for SoxN Ephrin Tl santa-maria
twf side pros dac
kay Glt Wnt4 eya
pum Rac2 sm Rab6 spri
sad sn NT1
RhoGEF64C kuz Gli how
fog Timp
NetA daw Trim9 Mp rk
Dg Rich Rab5 betaTub60D Ama
Mmp2 haf mid RhoL
NetB
rho zfh1 LRP1 Con
ham sens Sema2b
Dronc chic Sulf1 peb
Mtp Sdc if beat-IIa
pk Nrt beat-IIb
Nrk ena Tig
dar1
Wnt2 kn futsch
dati
Rop Manf
ss path
fax Optix
sick Sox14 hid
Spn mav
Mmp1
Sec23
Grip
wash kirre
NijC glec
Sgs3
fat-spondin
hbs m
Sgs4
Tsp
Sgs5 eater
Sgs5bis CG10339
Sgs8
Eig71Ee ItgaPS4
Cad88C
bib CG45263
CG30046 CG5758
NijA
Nlg1
Sgs7 Lac Sec24CD

**Figure S6: Gene ontology enrichment analysis of the genes downregulated in wing discs expressing *Gαq* under *C765-Gal4* driver (FC ≤ 0.65). A)** A dot plot displaying the top 25 GO terms that were significantly enriched among the statistically significantly downregulated genes (log2FC ≤ -0.6). The color of the dot represents the adjusted p-value, and the size represents the number of genes downregulated in the respective category. **B)** An emaplot showing the network representation of the top 25 significantly enriched GO terms grouped based on similarity. The color of the node represents the adjusted p-value, and the size represents the number of genes that were statistically significantly downregulated (log2FC ≤ -0.6). **C)** A cnet plot showing the top five enriched GO terms mapped to the corresponding genes which were statistically significantly upregulated. Each leaf node corresponds to a gene, and the node's color indicates the log fold change.

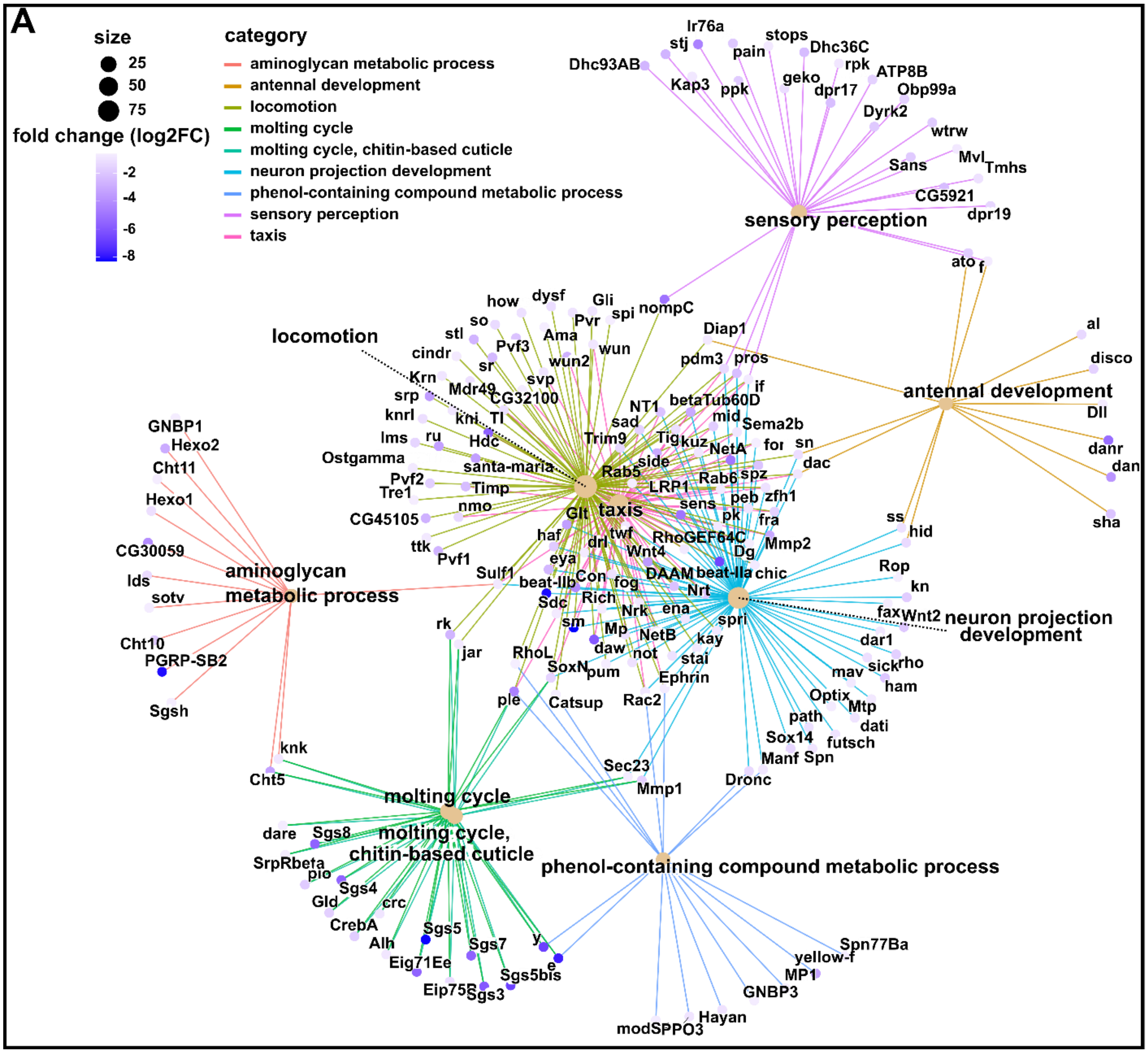


**Figure S7: Network representation of the top GO terms enriched among the significantly downregulated genes in wing discs expressing *Gαq* under *C765-Gal4* driver (log2FC ≤ -0.6). A)** A cnet plot depicting the top 10 enriched GO terms following the removal of redundant enriched GO terms and the corresponding genes that were statistically significantly downregulated. Each leaf node corresponds to a gene, and the node's color indicates the log fold change.

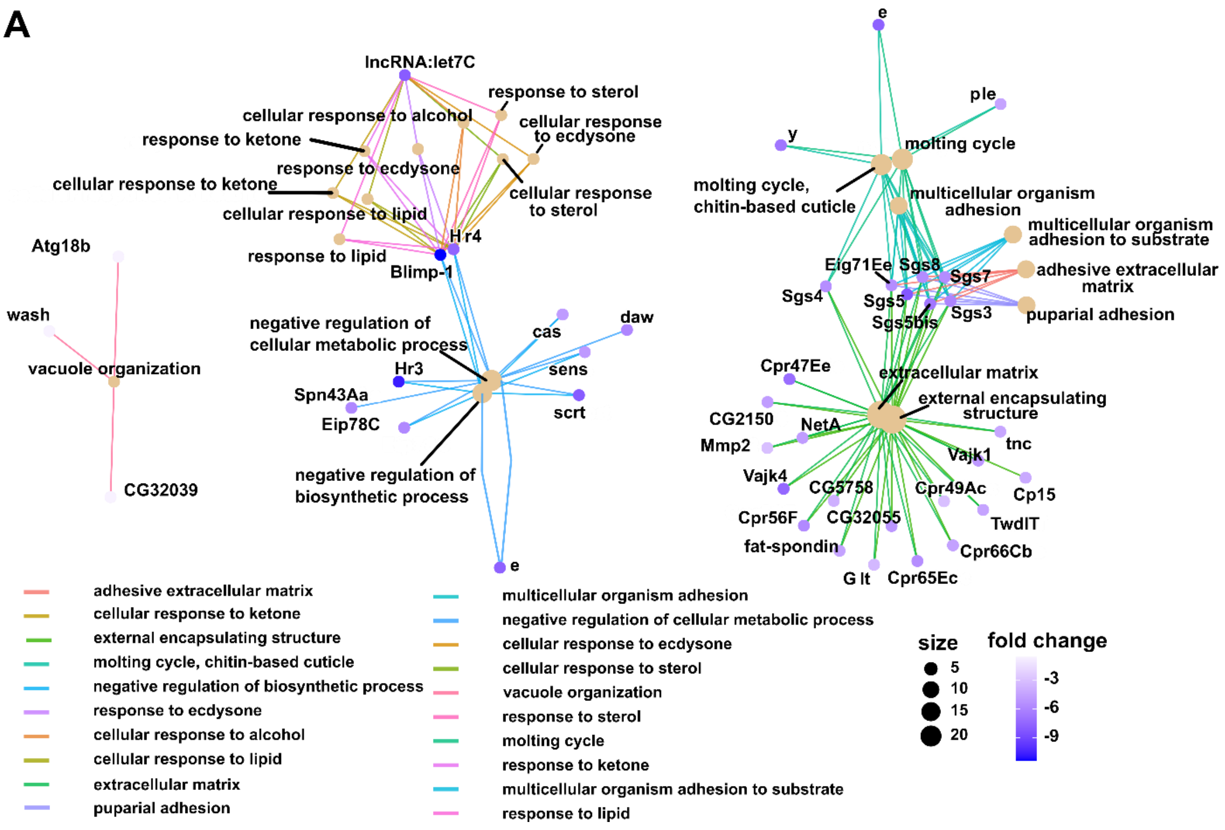


**Figure S8: Overexpression of *Gαq* in the wing disc under *C765* driver downregulated genes involved in the cellular response to ecdysone, extracellular matrix, and negative regulation of the metabolic process**. A) A cnet plot depicting the GO terms related to the molting cycle, cellular response to ecdysone, extracellular matrix, and the corresponding genes that were statistically significantly downregulated (log2FC ≤ -0.6). Each leaf node corresponds to a gene, and the node's color indicates the log fold change.

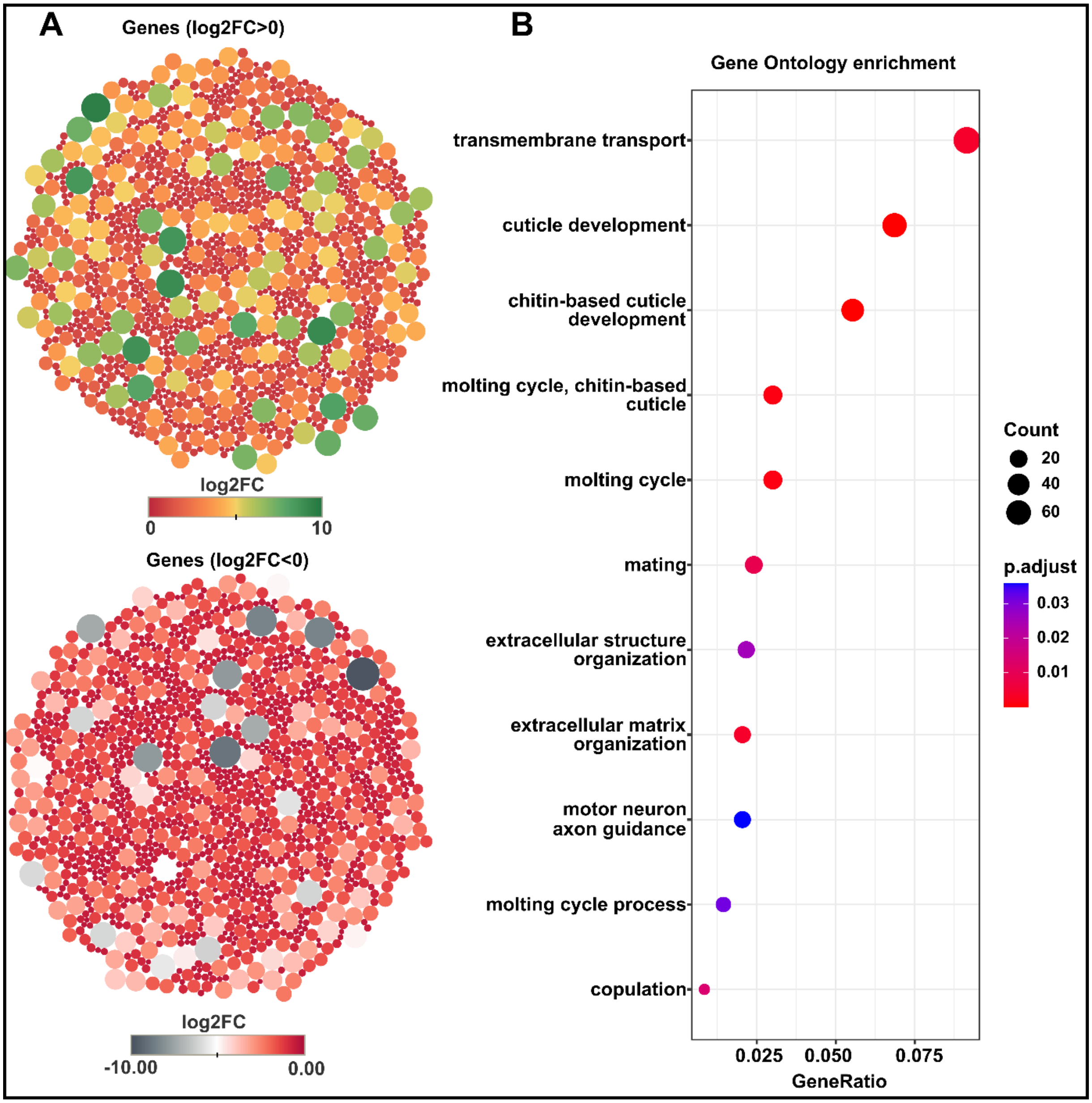


**Figure S9: Gene ontology enrichment analysis of differentially expressed genes in wing discs expressing *Gαq*$^{RNAi}$ (*C765>Gαq*$^{RNAi}$). A)** Bubble plot showing an overview of the genes that have a statistically significant fold change value in wing discs expressing *Gαq*$^{RNAi}$ under *C765-Gal4* driver. The top plot depicts the upregulated genes with a statistically significant fold change value, while the bottom plot depicts the downregulated genes with a statistically significant fold change value. Each bubble represents a gene. The bubble size represents the magnitude of the absolute log fold change value, whereas color represents the log fold change value. **B)** A dot plot displaying the top GO terms that were significantly enriched among the statistically significantly differentially expressed genes ($-0.6 \geq logFC \geq 0.6$). The gene ratio represents the ratio between the number of genes with statistically significant differential expression and the total number of genes representing the GO term.

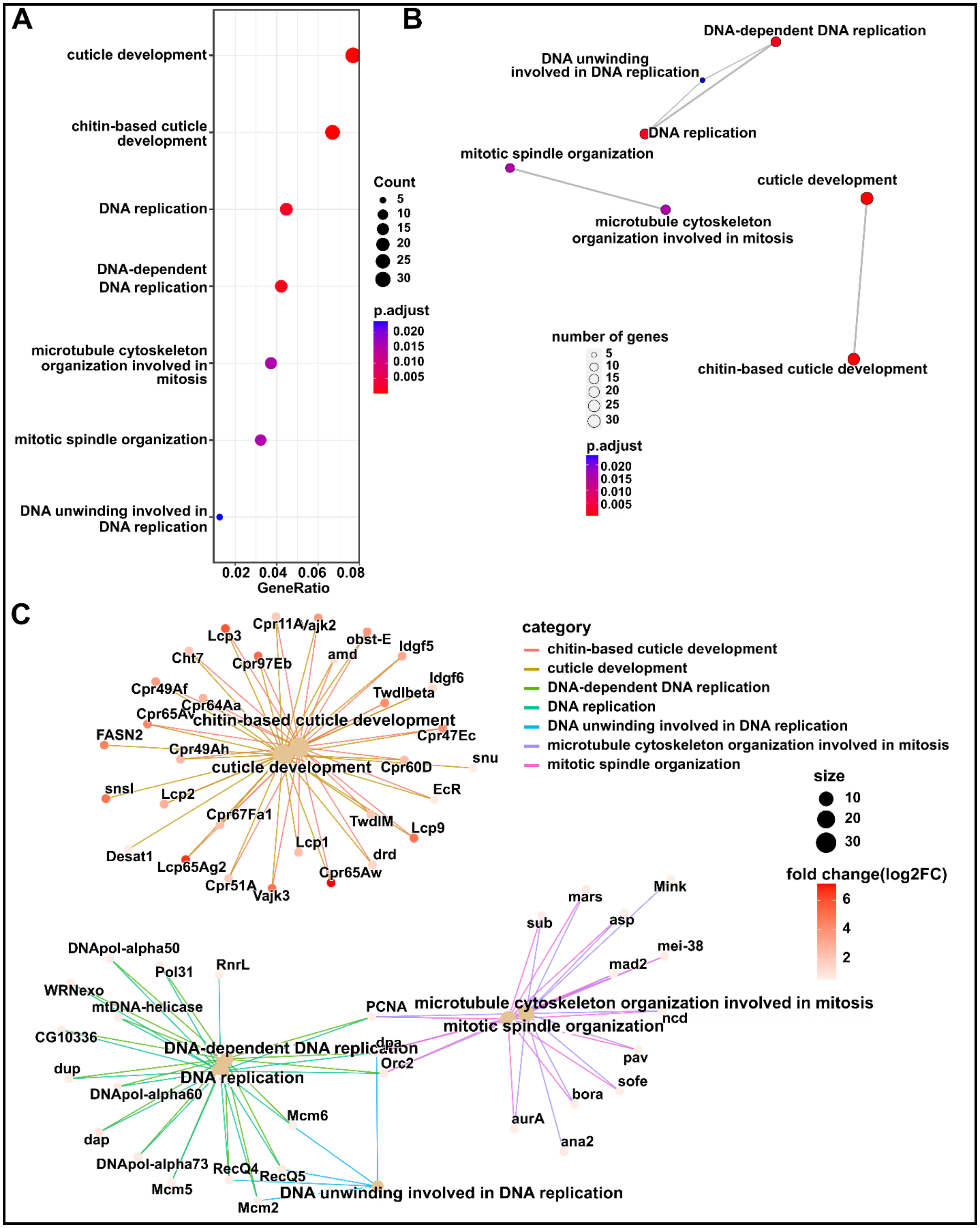
A
cuticle development
chitin-based cuticle development
DNA replication
DNA-dependent DNA replication
microtubule cytoskeleton organization involved in mitosis
mitotic spindle organization
DNA unwinding involved in DNA replication
0.02 0.04 0.06 0.08
GeneRatio
Count
5
10
15
20
25
30
p.adjust
0.020
0.015
0.010
0.005
B
DNA-dependent DNA replication
DNA unwinding involved in DNA replication
DNA replication
mitotic spindle organization
cuticle development
microtubule cytoskeleton organization involved in mitosis
chitin-based cuticle development
number of genes
5
10
15
20
25
30
p.adjust
0.020
0.015
0.010
0.005
C
category
chitin-based cuticle development
cuticle development
DNA-dependent DNA replication
DNA replication
DNA unwinding involved in DNA replication
microtubule cytoskeleton organization involved in mitosis
mitotic spindle organization
size
10
20
30
fold change(log2FC)
6
4
2
Lcp3
Cpr11A
Vajk2
obst-E
Idgf5
Cht7
amd
Cpr97Eb
Idgf6
Cpr49Af
Twdlbeta
Cpr64Aa
Cpr65Av
chitin-based cuticle development
FASN2
Cpr47Ec
Cpr49Ah
cuticle development
Cpr60D
snu
snsl
EcR
Lcp2
Cpr67Fa1
TwdlM
Lcp9
Lcp1
Desat1
drd
Lcp65Ag2
Cpr65Aw
Cpr51A
Vajk3
mars
Mink
sub
asp
mei-38
mad2
DNApol-alpha50
Pol31
RnrL
WRNexo
mtDNA-helicase
PCNA
microtubule cytoskeleton organization involved in mitosis
ncd
mitotic spindle organization
CG10336
dpa
DNA-dependent DNA replication
Orc2
pav
dup
DNA replication
sofe
DNApol-alpha60
bora
Mcm6
aurA
dap
ana2
DNApol-alpha73
RecQ4
RecQ5
Mcm5
DNA unwinding involved in DNA replication
Mcm2

**Figure S10: Gene Ontology enrichment analysis of upregulated genes for *$G\alpha q^{RNAi}$* expression in wing discs (*$C765>G\alpha q^{RNAi}$*) with log2FC ≥ 0.6. A)** A dot plot displaying the top GO terms that were significantly enriched among the statistically significantly upregulated genes (logFC ≥ 0.6). The color of the dot represents the adjusted p-value, and the size represents the number of genes upregulated in the respective category. **B)** Emaplot illustrating the network representation of the top significantly enriched GO terms that were grouped based on similarity. Each GO term is represented by a node. The color of the node represents the adjusted p-value, and size represents the number of genes that are statistically significantly upregulated in the GO term. **C)** A cnet plot showing the enriched GO terms mapped to the corresponding genes which were statistically significantly upregulated. Each leaf node corresponds to a gene, and the node's color indicates the log fold change.

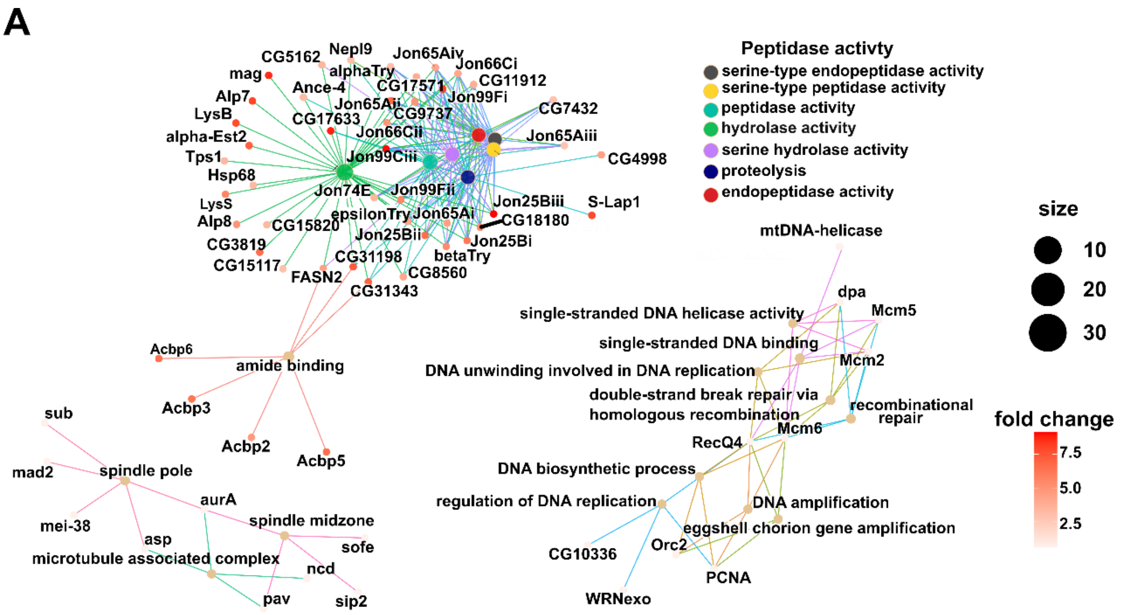


**Figure S11: Expression of *Gαq*$^{RNAi}$ in the wing disc under *C765* driver upregulated genes associated with peptidase activity and DNA replication**. A) A cnet plot showing network representation depicting the GO terms related to peptidase activity, endopeptidase activity, and the corresponding genes that were upregulated (log2FC ≥ 0.6). Each leaf node corresponds to a gene, and the node's color indicates the log fold change.

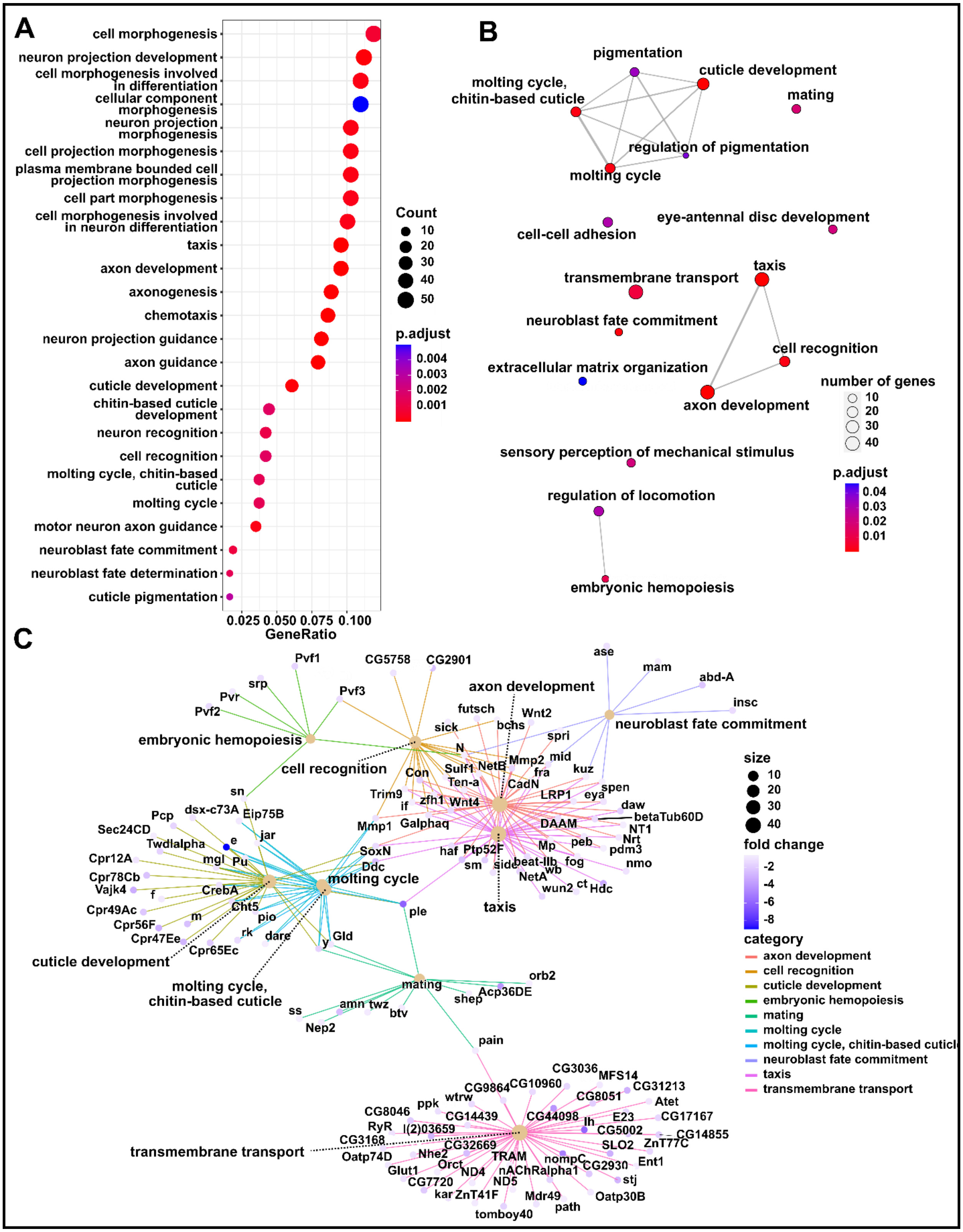
A
cell morphogenesis
neuron projection development
cell morphogenesis involved in differentiation
cellular component morphogenesis
neuron projection morphogenesis
cell projection morphogenesis
plasma membrane bounded cell projection morphogenesis
cell part morphogenesis
cell morphogenesis involved in neuron differentiation
taxis
axon development
axonogenesis
chemotaxis
neuron projection guidance
axon guidance
cuticle development
chitin-based cuticle development
neuron recognition
cell recognition
molting cycle, chitin-based cuticle
molting cycle
motor neuron axon guidance
neuroblast fate commitment
neuroblast fate determination
cuticle pigmentation
0.025 0.050 0.075 0.100
GeneRatio
Count
10
20
30
40
50
p.adjust
0.004
0.003
0.002
0.001
B
pigmentation
cuticle development
molting cycle, chitin-based cuticle
mating
regulation of pigmentation
molting cycle
cell-cell adhesion
eye-antennal disc development
transmembrane transport
taxis
neuroblast fate commitment
cell recognition
extracellular matrix organization
axon development
number of genes
10
20
30
40
sensory perception of mechanical stimulus
p.adjust
0.04
0.03
0.02
0.01
regulation of locomotion
embryonic hemopoiesis
C
axon development
embryonic hemopoiesis
cell recognition
neuroblast fate commitment
molting cycle
cuticle development
molting cycle, chitin-based cuticle
mating
taxis
transmembrane transport
size
10
20
30
40
fold change
-2
-4
-6
-8
category
axon development
cell recognition
cuticle development
embryonic hemopoiesis
mating
molting cycle
molting cycle, chitin-based cuticle
neuroblast fate commitment
taxis
transmembrane transport

**Figure S12: Gene Ontology enrichment analysis of downregulated genes for *Gαq*$^{RNAi}$ expression in wing discs (*C765>Gαq*$^{RNAi}$) with log2FC ≤ -0.6. A)** A dot plot displaying the top GO terms that were significantly enriched among the statistically significantly downregulated genes (FC ≤ 0.65). The color of the dot represents the adjusted p-value, and the size represents the number of genes downregulated in the respective category. **B)** An emaplot showing the network representation of the top significant enriched GO (redundant GO terms were removed to simplify) terms grouped based on similarity. The color of the node represents the adjusted p-value, and the size represents the number of genes that were statistically significantly downregulated (FC ≤ 0.65). **C)** Network representation of the top enriched GO terms following the removal of redundant GO terms and the corresponding genes that were statistically significant downregulated. Each leaf node corresponds to a gene, and the node's color indicates the log fold change.

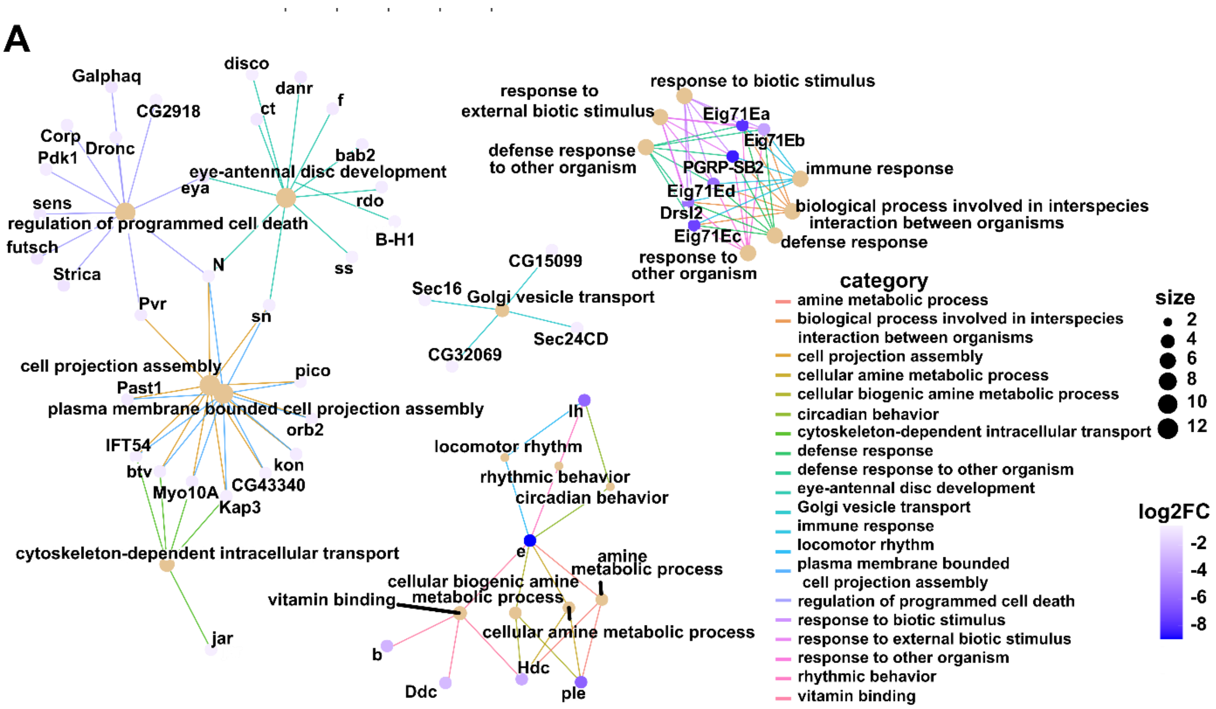


**Figure S13: Expression of *Gαq*$^{RNAi}$ in the wing disc driven by the *C765* driver downregulated genes involved in programmed cell death, ecdysone-induced genes response, and cell projection assembly**. A) A cnet plot showing network representation depicting the GO terms related to programmed cell death, cell projection assembly, defense response, immune response, and the corresponding genes that were downregulated. A leaf node corresponds to a gene whose log fold change is indicated by its color.

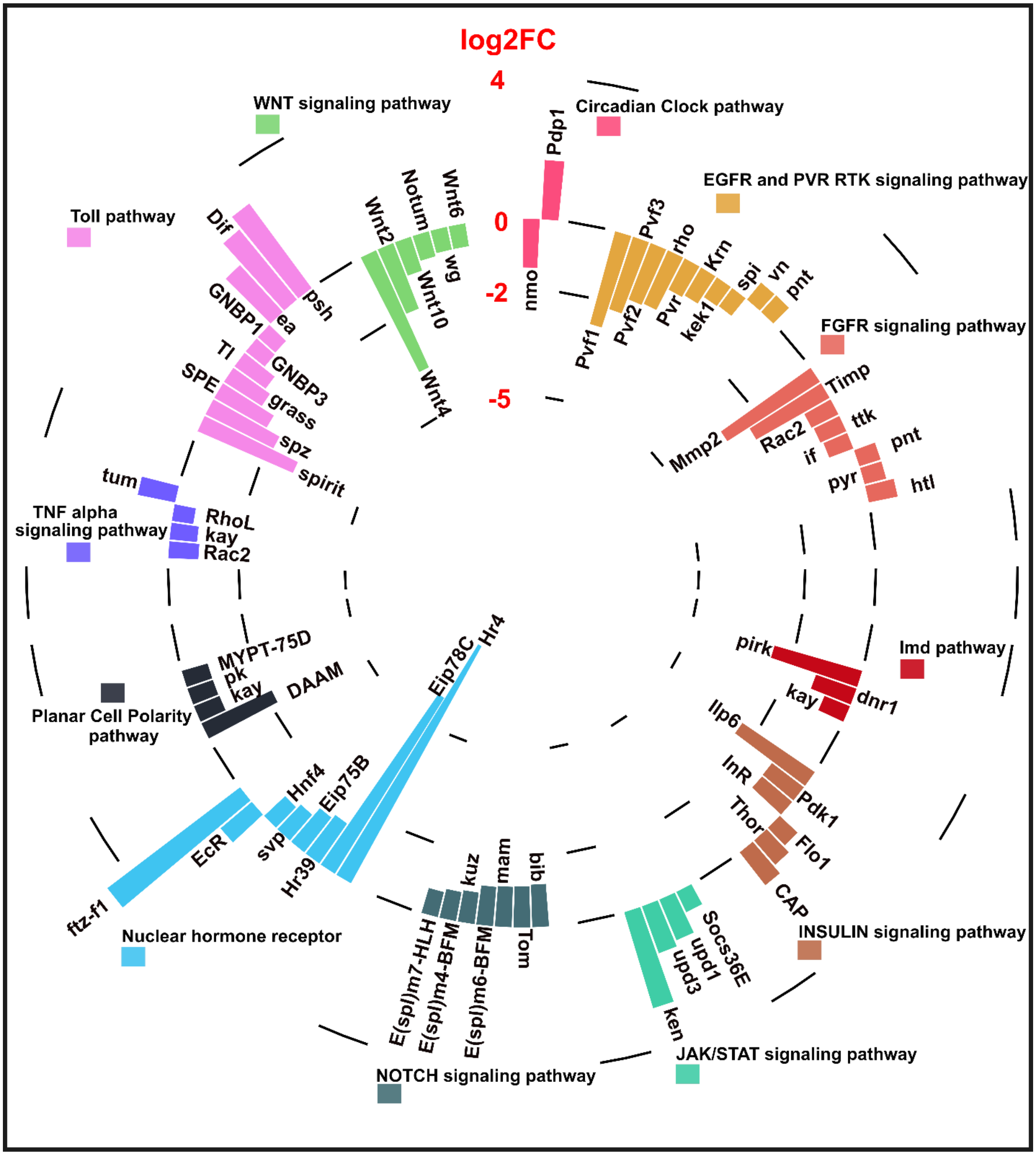


**Figure S14: Circular plot illustrating the fold change values of core signaling pathway genes that were differentially expressed in wing discs over expressing *Gαq* with the C765-Gal4 driver.**

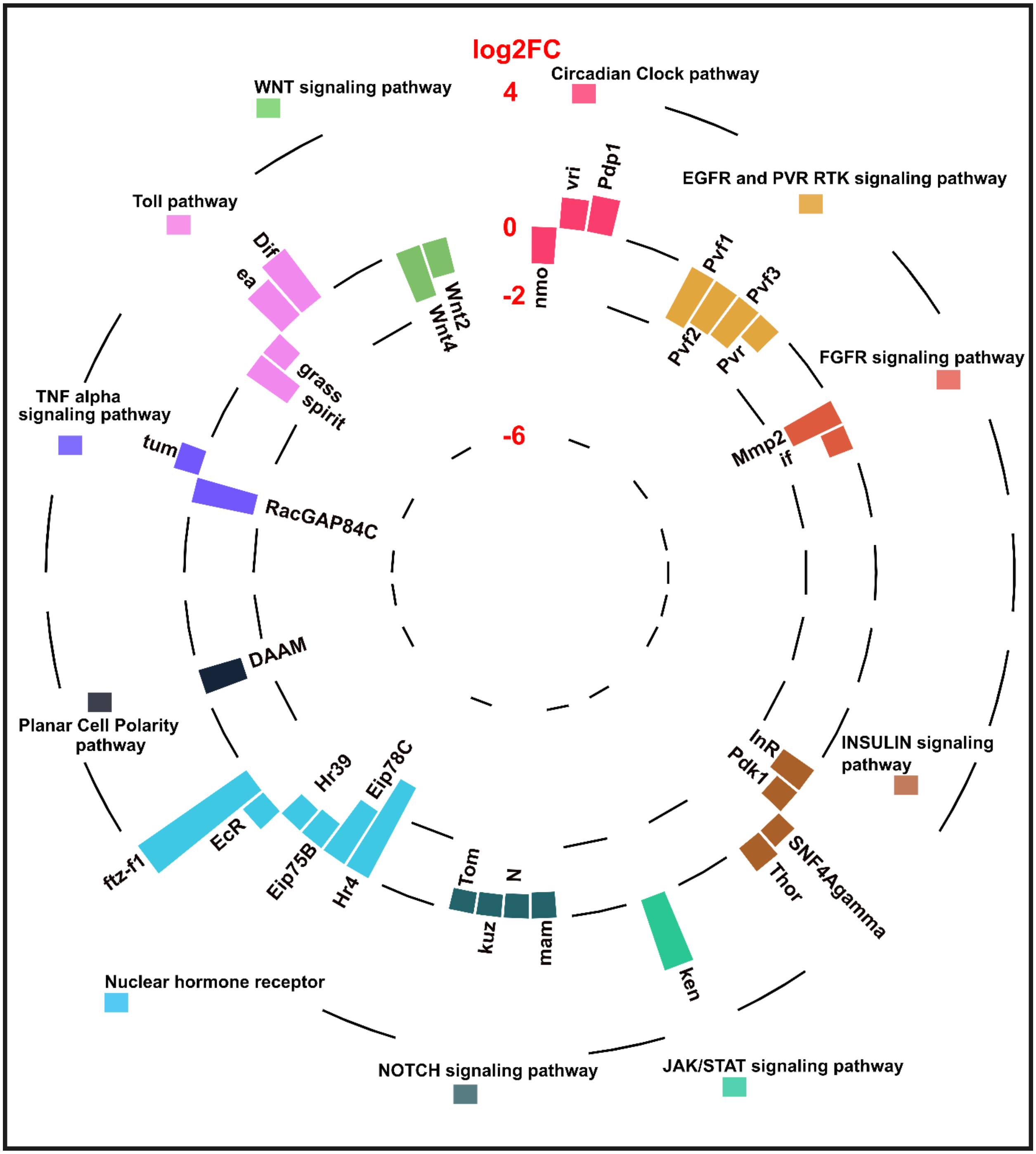


**Figure S15: Circular plot illustrating the fold change values of core signaling pathway genes that were differentially expressed in wing discs expressing *$G\alpha q^{RNAi}$* with the C765-Gal4 driver.**
